# Breath as an internal context: Respiratory phase alignment between encoding and retrieval optimizes memory performance

**DOI:** 10.64898/2025.12.25.696473

**Authors:** Nozomu H. Nakamura, Kohzoh Yoshino, Masaki Fukunaga

**Affiliations:** Department of Integrative Physiology, Faculty of Medicine, Hyogo Medical University, 1-1, Mukogawa cho, Nishinomiya, Hyogo 663-8501 Japan; Department of Biomedical Sciences, School of Biological and Environmental Sciences, Kwansei Gakuin University, 1, Gakuen Uegahara, Sanda, Hyogo 669-1330 Japan; Section of Brain Function Information, National Institute for Physiological Sciences, 38 Nishigonaka Myodaiji, Okazaki, Aichi 444-8585, Japan; Core for Spin Life Sciences, Okazaki Collaborative Platform, National Institutes of Natural Sciences, 38 Nishigonaka Myodaiji, Okazaki 444-8585, Japan; Graduate Institute for Advanced Studies, SOKENDAI, Hayama, Kanagawa 340-0193, Japan

**Author notes:** **Corresponding Author:** Nozomu H. Nakamura, Department of Integrative Physiology, Faculty of Medicine, Hyogo Medical University, 1-1, Mukogawa cho, Nishinomiya, Hyogo 663-8501 Japan.

**Keywords:** encoding, retrieval, inspiratory onset, phase matching

## Abstract

Increasing evidence suggests that respiratory rhythms are crucial for cognitive functions. Although recent findings suggest that respiratory activity may serve as an internal contextual framework during memory processes, the contribution of specific respiratory phases to encoding and retrieval remains unclear. Here, we investigated the breathing-dependent performance of 30 healthy volunteers during a visual delayed matching-to-sample recognition memory task while their nasal respiration was monitored. Reaction times (RTs) decreased when visual cues were both encoded and retrieved during the late phase of exhalation. In contrast, longer RTs were observed during late exhalation at retrieval when encoding occurred either (i) during a period that encompassed inspiratory onset (i.e., exhalation-to-inhalation transition or EI transition) or (ii) during the early phase of inhalation. These contrasting outcomes under identical retrieval conditions highlight the phase-dependent effects of respiration, specifically the alignment of respiratory timing between encoding and retrieval. Importantly, we found that the late phase of exhalation during both encoding and retrieval may represent a favorable temporal window for shaping memory performance. These findings suggest that respiratory phase alignment modulates memory processes by providing an interoceptive, phase-dependent context for cognition.

## Introduction

Memory encoding and retrieval are inherently embedded within contextual frameworks[1,2]. Context is the spatiotemporal informational background that integrates external environmental states with internal bodily states[3,4]. Considerable evidence suggests that interoceptive signals, such as respiration and cardiac pulses, may function as internal contextual cues that scaffold and modulate cognitive function[5–7].

Interoception refers to the brain’s perception and integration of internal bodily signals[8]. It enables the body to maintain stability while adapting to changing demands ranging from homeostasis to allostasis[9,10]. Interoceptive signals are continuously fluctuating and are considered “dynamic internal contexts” capable of shaping information processing in the brain. Human studies have demonstrated that respiratory dynamics modulate a wide range of cognitive functions, including neural excitability, perception, memory, and motor actions[11–16]. Moreover, the timing and coordination of respiratory phases influence activity across limbic and attentional networks and have been linked to moment-to-moment variations in cognitive performance[17–20]. Our previous work further demonstrated that the inclusion of inspiratory onset – i.e., the exhalation-to-inhalation (EI) transition – during retrieval is associated with reduced accuracy and prolonged reaction times (RTs) [21,22]. This effect was likely driven by either the use of a 6-sec task (test) cycle (a response window of 6 sec between cue exposure and pressing a button)[22] or respiratory phase locking during the task (3.5 sec or more in the human respiratory cycle)[21], which allowed alignment with the human respiratory rhythm. Importantly, this effect diminished substantially when the test cycle was every 3 sec or less than 3 sec[21], indicating that test-cycle alignment with respiration at retrieval is critical for such EI-transition-dependent effects.

Notably, our interventional study using genetically modified mice and optogenetics revealed that inspiratory-onset (or EI-transition) activity precisely timed to encoding modulates memory performance[23]. In this study, we selectively manipulated inspiratory-onset signals originating from the PreBötzinger complex (PreBötC) in the brainstem – the primary inspiratory rhythm generator[24–27] – and observed both enhancement and impairment of memory performance mediated by hippocampal ensemble dynamics[23]. These findings suggest that the timing of respiration may play a crucial role in creating a temporal scaffold for memory formation[23,28]. The key element of the respiratory phase is thought to be the alignment of context-related information between encoding and retrieval. In this framework, it remains unclear whether the inhalation and exhalation phases can constitute a “reinstatable internal context” such that memory performance is optimized when encoding and retrieval occur within matching respiratory phases.

Here, we investigated whether phase-dependent alignment of respiration between encoding and retrieval modulates task performance (i.e., RT and accuracy). To rigorously isolate encoding-dependent effects, we employed a shorter version of a delayed matching-to-sample (DMTS) task with a 2-sec test cycle. The use of a test cycle of less than 3 sec prevents EI-transition-dependent cognitive decline during retrieval, as previously reported[21], thereby enabling a clearer assessment of respiration-related influences during encoding. We applied an autoregressive moving average (ARMA) model, which can remove history-dependent fluctuations from the time series of cognitive parameters to disentangle intrinsic temporal dynamics in task performance across the respiratory cycle. Analysis of the residual fluctuations in task performance revealed a phase alignment of respiration, suggesting that respiratory activity provides a phase-dependent internal context modulating memory performance.

## Results

### Memory performance and respiratory dynamics

In the present study, 30 healthy volunteers performed a visual DMTS task, which required the memorization and recognition of natural objects[21,22] in the photographs (Fig. 1a) of the THINGS database[29,30]. Each participant memorized 40 photographs (sample cues) and discriminated 80 photographs (test cues) in each session. In total, individual participants performed 400 memorizations and 800 discriminations. In the sample block, each sample cue was continuously presented on the screen every 1 sec, and each sample event lasted 1 sec (Fig. 1b). In the test block, a test event or RT was defined as the period from test cue presentation to a button-pressed response. With respect to the designated DMTS task, although RT gradually increased and total accuracy (accuracy of test events) and hit scores (accuracy of ‘old’ events) gradually decreased within a session, these cognitive parameters did not differ across sessions (Suppl. Fig. S1). These results indicate that the volunteers maintained a comparable level of task performance throughout the sessions. Notably, there was no difference in the correct rejection (CR) scores (accuracy of ‘new’ events) within the session or among the sessions.

**Fig. 1.**
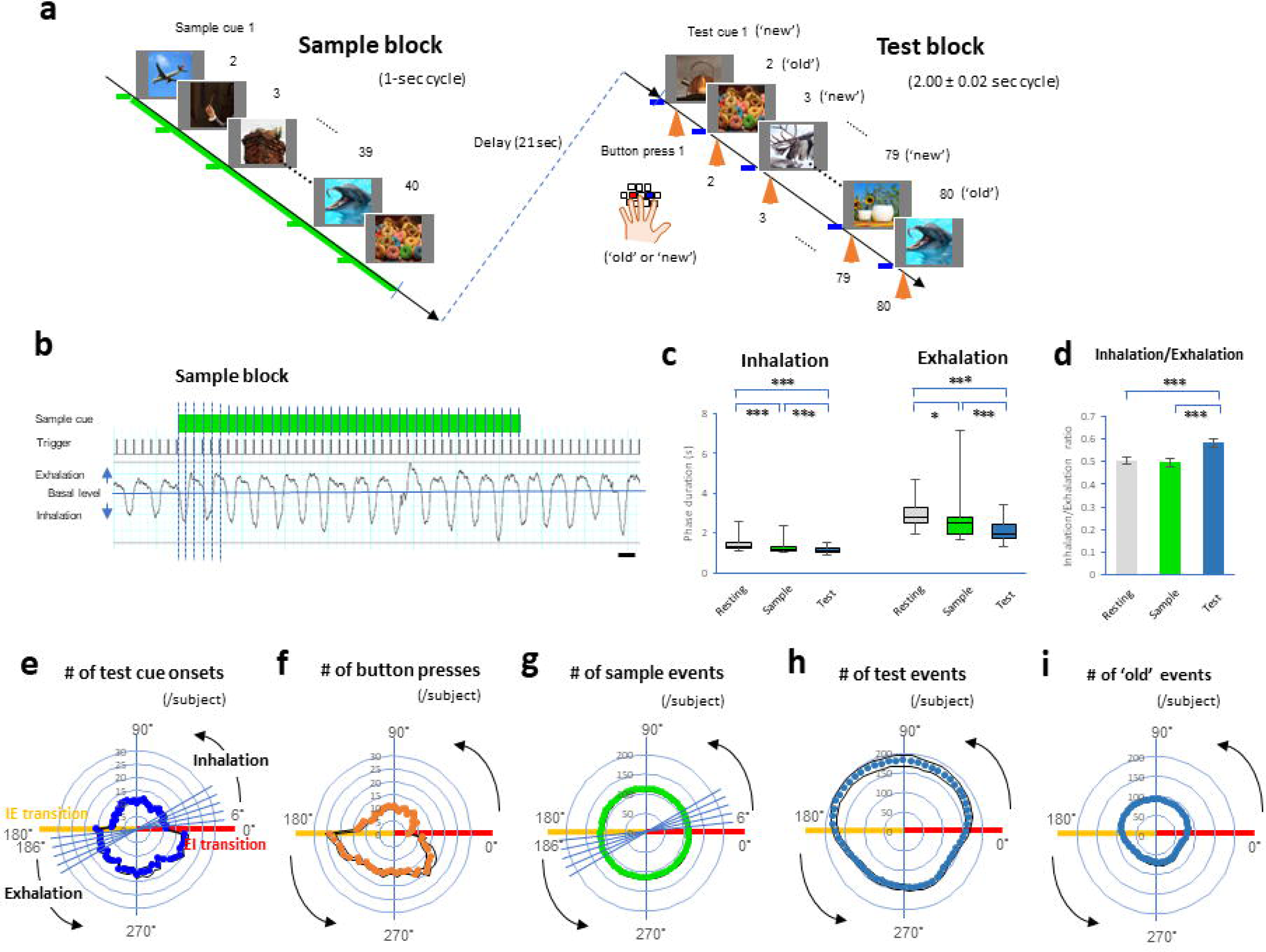
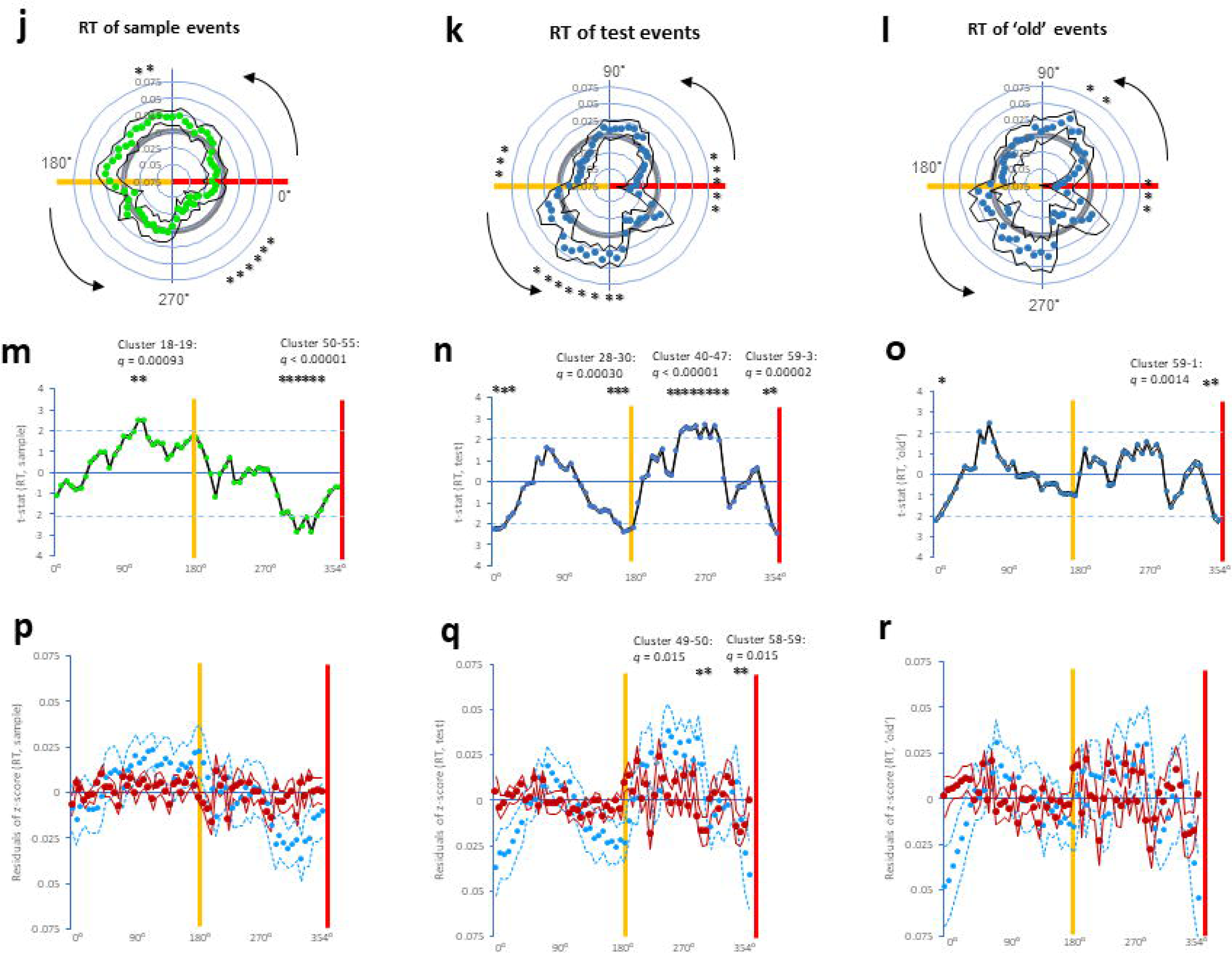
The delayed matching-to-sample (DMTS) task, circular distribution of the respiratory phase, and respiration-timing-dependent RT variations. **a.** DMTS task paradigm consisting of a sample block containing 40 sample cues (green), a delay block, and a test block containing 80 test cues (blue). Button presses (orange arrows) were based on the observed decisions. b. Representative plots showing the timing of 40 sample cues (green) and triggers along with a respiratory waveform during the sample block. Each sample cue (green) was continuously presented on the screen every 1 sec, and each sample event lasted 1 sec. c, d. Plots showing durations of inhalation (left panel in c), exhalation (right panel in c), and the ratio of inhalation to exhalation (d), among the resting state (gray), the sample block (green), and the test block (blue). e, f. Plots showing circular histograms of test cue onset (blue, e) and button-press response (orange, f) in the respiratory cycle. The cognitive data were plotted in each 6° bin in a total of 60 bins (30 bins during inhalation: 0-180°; 30 bins during exhalation: 180-360°). g-i. Plots showing circular histograms of sample events (green, g), test events (dark blue, h), and ‘old’ events (dark blue, i), where the overlap of individual events with each bin was counted throughout the respiratory cycle. j-l. Plots showing circular distributions of z-scored RT of sample events (j), test events (k), and ‘old’ events (l) in the respiratory cycle. m-o. Plots showing t-statistics of z-scored RTs of sample events (m), test events (n), and ‘old’ events (o) together with cluster formations. p-r. Plots showing residuals (dark red) of z-scored RTs of sample events (p), test events (q), and ‘old’ events (r) together with cluster formations. The light blue dots indicate the ARMA (1, 1) fitted dots. The scale bar in b indicates 2 sec. The yellow and red lines in e-r indicate the IE transition (exhalation onset) and EI transition (inhalation onset), respectively. * *p* ≤ 0.05 in c, d, and j-l, *** *p* ≤ 0.005 in c and d, ** *q* ≤ 0.05 (a single cluster with two bins), *** *q* ≤ 0.05 (a single cluster with three bins), and ***** *q* ≤ 0.05 (a single cluster with five bins) in m, n, o, and q.

With respect to respiratory dynamics, the Shapiro-Wilk normality test did not reveal a normal distribution of durations of inhalation or exhalation (Fig. 1c). There were significant differences among the sample blocks, test blocks, and resting blocks before and after the tasks (5 min each) in terms of the durations of inhalation (χ^2^(2) = 37.267, *p* = 8.1 × 10^−9^, Friedman test, left panel in Fig. 1c), exhalation (χ^2^(2) = 41.267, *p* = 1.1 × 10^−9^, right panel in Fig. 1c), and respiration (inhalation + exhalation, Suppl. Fig. S2). Post hoc comparisons revealed that the test block presented the shortest durations of inhalation and exhalation, and the sample block presented the second shortest durations of inhalation and exhalation (Wilcoxon signed-rank test with Bonferroni correction). Interestingly, the ratio of inhalation to exhalation was greater in test blocks than in resting and sample blocks (*F*(2, 58) = 16.5, *p* = 2.1 × 10^−6^, one-way repeated-measures ANOVA; test block: *p* = 0.0005 to resting block, *p* = 5.7 × 10^−7^ to sample block, post hoc pairwise comparisons via paired *t* tests with the Bonferroni correction, Fig. 1d), indicating that exhalation was shortened during retrieval.

### Respiration-timing-dependent cognitive variations

Averaged cognitive parameters across individuals were plotted in 60 angular bins of 6° each (inhalation: 0-180°; exhalation: 180-360°), with one full circle representing a complete respiratory cycle. Histograms of test cue onset (Fig. 1e) and button-press responses (Fig. 1f) across the respiratory cycle exhibited skewed distributions. We also constructed circular histograms of sample events, test events, and ‘old’ events (matching events, Fig. 1g-i). The duration of sample events was fixed at 1 s, whereas the durations of test and ‘old’ events – defined as the interval from cue exposure to the button press – were approximately 0.6 s. Overlapping portions of sample, test, and ‘old’ events were counted in each angular bin. To evaluate respiration-timing-dependent effects on task performance, individual parameters of sample events, test events, and ‘old’ events in each of the 60 bins of the respiratory cycle were used for subsequent analysis.

We carried out respiratory phase (bin)-dependent analysis of RT and the accuracy of sample events, test events, and ‘old’ events (see Methods). We found a skewed distribution of z-scored RTs of sample events throughout the sample respiratory cycle (*ps* ≤ 0.05, two-tailed one sample *t*-test against zero, Fig. 1j), z-scored RTs of test events during the test respiratory cycle (*ps* ≤ 0.05, Fig. 1k), and z-scored RTs of ‘old’ events during the test respiratory cycle (*ps* ≤ 0.05, Fig. 1l). The nonparametric permutation test and false discovery rate (FDR) correction revealed the significance of cluster formations (more than two adjacent bins) in the RT of sample events (cluster 18-19: *t*_(sum)_ = 5.02, *q* = 0.00093; cluster 50-55: *t*_(sum)_ = 14.65, *q* < 1.0 × 10^−5^, Fig. 1m), RT of test events (cluster 28-30: *t*_(sum)_ = 6.88, *q* = 3.0 × 10^−4^; cluster 40-47: *t*_(sum)_ = 19.80, *q* < 1.0 × 10^−5^; cluster 59-3: *t*_(sum)_ = 11.07, *q* = 1.5 × 10^−5^, Fig. 1n), and RT of ‘old’ events (cluster 59-1: *t*_(sum)_ = 6.41, *q* = 0.0014, Fig. 1o). Moreover, ARMA (*p* = 1, *q* = 1) modeling and residual analysis via a permutation test and FDR correction revealed that the residuals formed clusters in the RTs of the test events (cluster 49-50: *t*_(sum)_ = 4.38, *q* = 0.015; cluster 58-59: *t*_(sum)_ = 4.35, *q* = 0.015, Fig. 1q). There were also significant cluster formations in terms of total accuracy (accuracy of the test events) and hit score (accuracy of the ‘old’ events), whereas the accuracy of sample events did not have any cluster formation (Suppl. Fig. S3). However, ARMA (1, 1) modeling and residual analysis did not prove residual cluster formation in terms of total accuracy or hit score, even though accuracy tended to increase during the early phase of inhalation.

Residual analysis revealed cluster formations (more than two adjacent bins) among the RT of test events during late exhalation (Fig. 1q). Since the residuals obtained from the ARMA (1, 1) model reflect binwise deviations that are free from serial dependencies, the consistency of cluster formations of residuals at the adjacent or same bins (i.e., cluster 49-50, cluster 58-59 in Fig. 1q) can be interpreted as reflecting respiration-timing-dependent RT variations that are independent of autocorrelated noise.

### Effects of respiratory conditions

In line with our previous study[22], test events were classified into the following four respiratory conditions (Fig. 2a and 2b): (i) the test NoI (NoI_t_) condition (staying within the inhalation without the transitions; see Methods); (ii) the test IEt (IEt_t_) condition (containing the inhalation-to-exhalation transition or IE transition); (iii) the test NoE (NoE_t_) condition (staying within the exhalation without the transitions); and (iv) the test EIt (EIt_t_) condition (containing inspiratory onset or the EI transition). The RT and accuracy of individual test events were averaged within each volunteer for each condition and were not normally distributed. We detected significant differences in the RT of test events (χ^2^(3) = 77, *p* < 2.2 × 10^−16^, Friedman test, Fig. 2c), the total accuracy of test events (χ^2^(3) = 14.40, *p* = 0.0024, Fig. 2d), the RT of ‘old’ events (*^2^*(3) = 64.72, *p* = 5.8 × 10^−14^, Fig. 2E), and the hit score (χ^2^(3) = 9.88, *p* = 0.020, Fig. 2f). No significant difference in the CR score was observed (Suppl. Fig. S4). Post hoc comparisons revealed that both the IEt_t_ and EIt_t_ conditions presented the highest RTs for test events and ‘old’ events, whereas the NoI_t_ condition presented the lowest RTs for test events and ‘old’ events (Wilcoxon signed rank test with Bonferroni correction). On the other hand, there was no clear difference in the accuracy of test events or ‘old’ events (hit scores) across the conditions, except that the IEt_t_ condition had a lower RT than the NoI_t_ condition did.

**Fig. 2.**
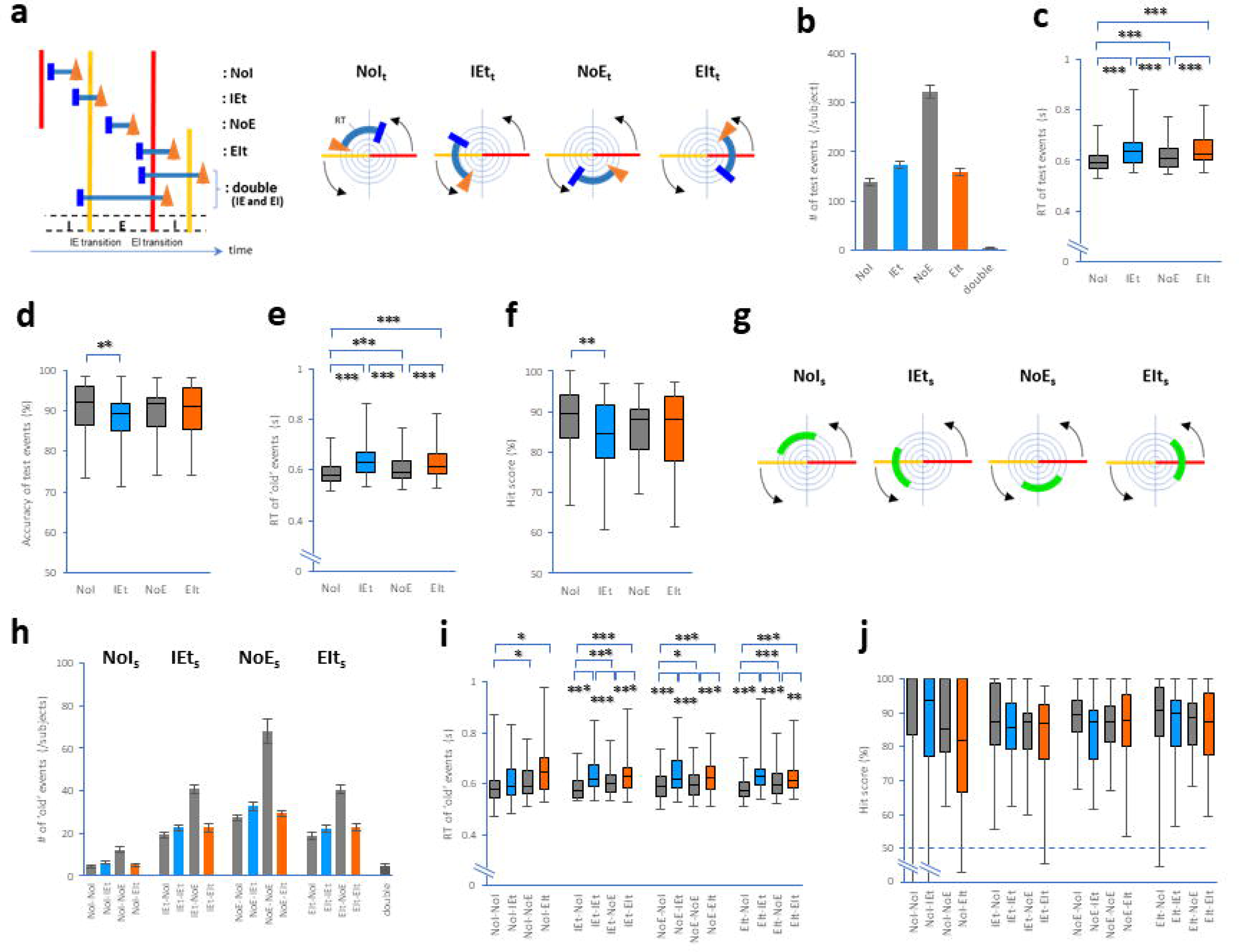
Effects of respiratory conditions. **a.** Drawing showing conditions of combinations between test events and transitions/phases of the respiratory cycle: (i) test events that occurred within the inhalation (NoI_t_ condition); (ii) test events that encompassed the IE transition (IEt_t_ condition); (iii) test events that occurred within the exhalation (NoE_t_ condition); (iv) test events that encompassed the EI transition (EIt_t_ condition); and (v) test events that encompassed both IE and EI transitions (a condition designated ‘double’). **b-f**. Plots showing the number of test events (**b**), RT of test events (**c**), accuracy of test events (**d**), RT of ‘old’ events (**e**), and hit score (accuracy of ‘old’ events, **f**). **g**. Drawing showing conditions of combinations between sample events and transitions/phases of the respiratory cycle: (i) sample events that occurred within the inhalation (NoI_s_ condition); (ii) sample events that encompassed the IE transition (IEt_s_ condition); (iii) sample events that occurred within the exhalation (NoE_s_ condition); (iv) sample events that encompassed the EI transition (EIt_s_ condition); and (v) sample events that encompassed both IE and EI transitions (a condition designated ‘double’). **h-j**. Plots showing the number of ‘old’ events (**h**), RT of ‘old’ events (**i**), and hit score (**j**). * *p* ≤ 0.05, ** *p* ≤ 0.01, and *** *p* ≤ 0.005.

According to the classification of sample events into four conditions (Fig. 2g, Suppl. Fig. S5), we divided the test events into the following conditions (Fig. 2h): the sample NoI (NoI_s_) series: (i) NoI_s_-NoI_t_, (ii) NoI_s_-IEt_t_, (iii) NoI_s_-NoE_t_, and (iv) NoI_s_-EIt_t_ conditions; the sample IEt (IEt_s_) series: (v) IEt_s_-NoI_t_, (vi) IEt_s_-IEt_t_, (vii) IEt_s_-NoE_t_, and (viii) IEt_s_-EIt_t_ conditions; the sample NoE (NoE_s_) series: (ix) NoE_s_-NoI_t_, (x) NoE_s_-IEt_t_, (xi) NoE_s_-NoE_t_, and (xii) NoE_s_-EIt_t_ conditions; and the sample EIt (EIt_s_) series: (xiii) EIt_s_-NoI_t_, (ivx) EIt_s_-IEt_t_, (vx) EIt_s_-NoE_t_, and (vix) EIt_s_-EIt_t_ conditions. RT and hit scores were not normally distributed. There were significant differences in the RTs of the sample respiratory condition series (NoI_s_ series: χ^2^(3) = 13.84, *p* = 0.0031; IEt_s_ series: χ^2^ (3) = 29.56, *p* = 1.7 × 10 ; NoE_s_ series: χ^2^ (3) = 48.60, *p* = 1.6 × 10 ; EIt_s_ series: χ^2^ (3) = 55.36, *p* = 5.8 × 10^−12^, Friedman test, Fig. 2i). Post hoc comparisons revealed that in the IEt_s_, NoE_s_, and EIt_s_ series, both the IEt_t_ and EIt_t_ conditions presented the highest RTs, whereas the NoI_t_ condition presented the lowest RT (Wilcoxon signed rank test with Bonferroni correction). No differences were observed in the hit scores (NoI_s_ series: χ^2^(3) = 3.84, *p* = 0.28; IEt_s_ series: χ^2^(3) = 4.89, *p* = 0.18; EIt_s_ series: _χ_*^2^*(3) = 0.983, *p* = 0.81, Fig. 2j), except in the NoE_s_ series (χ*^2^*(3) = 9.11, *p* = 0.028). However, post hoc comparison did not reveal any difference in the NoE_s_ series. Regardless of sample respiratory conditions, RT was primarily increased in the IEt_t_ and EIt_t_ conditions, whereas RT was preferentially decreased in the NoI_t_ condition.

### Circular respiratory analysis with phase conditions

The detailed mechanisms underlying encoding-induced coordination remain unclear. Here, we conducted a respiration-timing-dependent analysis of RT based on the following classification of sample conditions. We divided the datasets into four sample respiratory conditions (NoI_s_, IEt_s_, NoE_s_, and EIt_s_ conditions) and found skewed distributions of the z-score RT of the NoI_s_ (*ps* ≤ 0.05, two-tailed one sample *t*-test against zero, Fig. 3a), RT of the IEt_s_ (*ps* ≤ 0.05, Fig. 3b), RT of the NoE_s_ (*ps* ≤ 0.05, Fig. 3c), and RT of the EIt_s_ (*ps* ≤ 0.05, Fig. 3d) conditions during the test respiratory cycle. After a one-sample *t* test against zero in the individual bins, a nonparametric permutation test and FDR correction revealed the significance of cluster formations (more than two adjacent bins) in the RT of the NoI_s_ (cluster 39-40: *t*_(sum)_ = 4.87, *q* = 0.0010, Fig. 3a_2_), IEt_s_ (cluster 29-30: *t*_(sum)_ = 4.64, *q* = 0.0021, Fig. 3b_2_), and EIt_s_ (cluster 55-56: *t*_(sum)_ = 5.43, *q* = 0.00098, Fig. 3d_2_) conditions. ARMA (1, 1) modeling and residual analysis via a permutation test and FDR correction revealed that the residuals formed clusters in the RT of the EIt_s_ condition (cluster 57-58: *t*_(sum)_ = 5.21, *q* = 0.0013, Fig. 3d_3_). These results revealed that the sample conditions containing the EI transition (inspiratory onset; EIt_s_ condition) increased the RT during the late phase of exhalation.

**Fig. 3.**
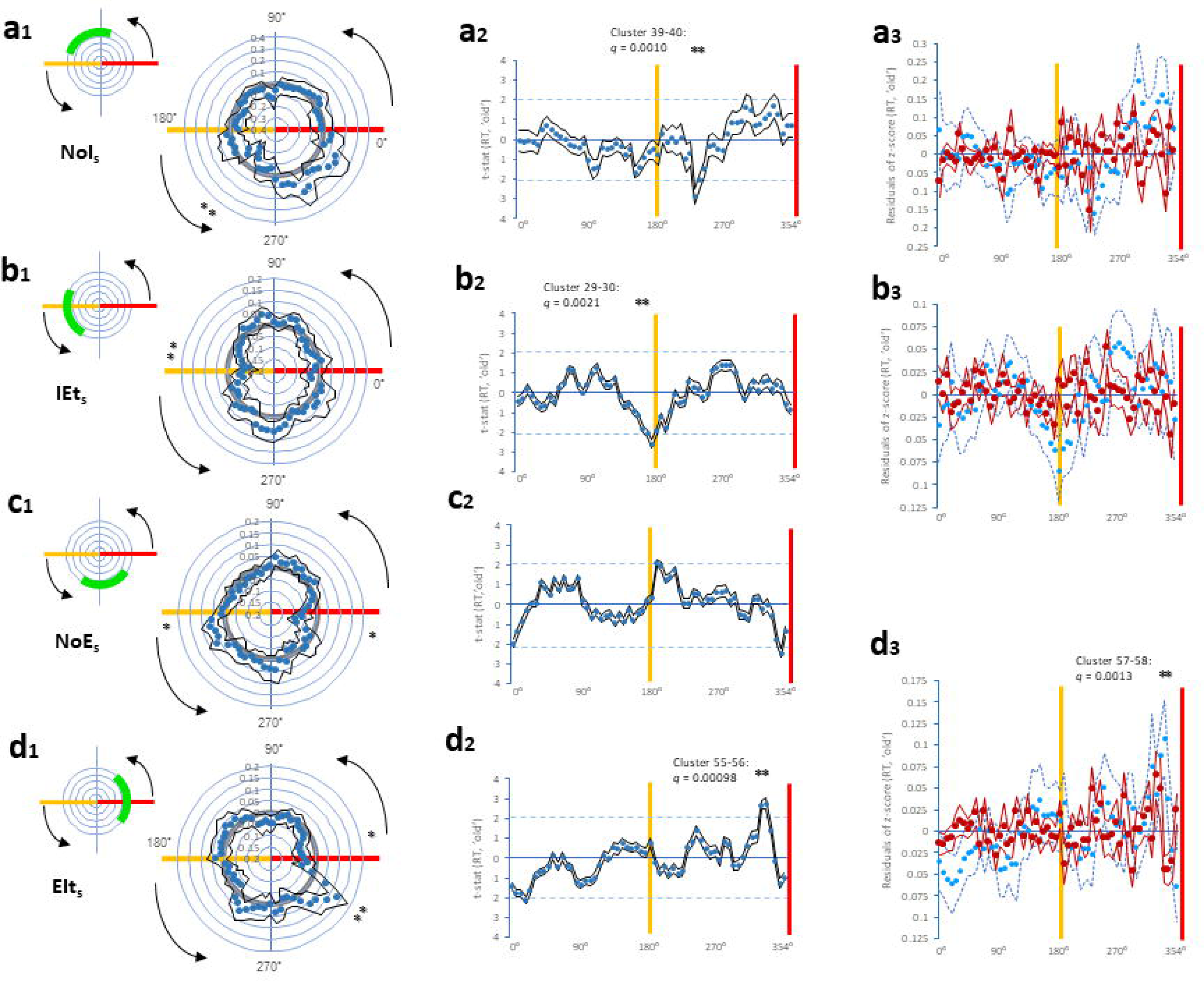
Cluster formations in RT of respiratory phase matching between sample and test blocks (1) Plots showing z-scored RTs (**a_1_**, **b_1_**, **c_1_**, **d_1_**), t-statistics (**a_2_**, **b_2_**, **c_2_**, **d_2_**), and residuals of z-scored RTs (**a_3_**, **b_3_**, **d_3_**) of ‘old’ events during the test respiratory cycle, according to four sample respiratory conditions: NoI_s_ condition (**a_1_-a_3_**), IEt_s_ condition (**b_1_-b_3_**), NoE_s_ condition (**c_1_, c_2_**), and EIt_s_ condition (**d_1_-d_3_**). * *p* ≤ 0.05 in **a_1_**, **b_1_**, **c_1_**, and **d_1_**. ** *q* ≤ 0.05 (a single cluster with two bins) in **a_2_**, **b_2_**, **d_2_** and **d_3_**.

To further elucidate the critical differences between the effects under and over the EI and IE transitions, we divided the RT dataset of sample events into the following four conditions: (i) RT of the sample In1 (the overlap of sample events in the 0-90° bin, Fig. 4a), (ii) RT of the sample In2 (the overlap of sample events in the 90-180° bin, Fig. 4b), (iii) RT of the sample Ex1 (the overlap of sample events in the 180-270° bin, Fig. 4c), and (iv) RT of the sample Ex2 (the overlap of sample events in the 270-360° bin, Fig. 4d) conditions. Their distributions were plotted along the test respiratory cycle. The nonparametric permutation test and FDR correction revealed the significance of cluster formation in the RT of the sample In1 (cluster 15-16: *t*_(sum)_ = 4.38, *q* = 0.0044; cluster 54-57: *t*_(sum)_ = 11.40, *q* < 1.0 × 10^−5^, Fig. 4a_2_), and sample Ex2 (cluster 59-3: *t*_(sum)_ = 12.02, *q* < 1.0 × 10^−5^, Fig. 4d_2_) conditions. ARMA (1, 1) modeling and residual analysis via a permutation test and FDR correction revealed that the residuals formed clusters in the RT of the sample In1 (cluster 54-55: *t*_(sum)_ = 4.15, *q* = 0.013, Fig. 4a_3_) and sample Ex2 (cluster 57-59: *t*_(sum)_ = 6.98, *q* = 4.0 × 10^−4^, Fig. 4d_3_) conditions. These results indicated that learning sample cues during the early phase of inhalation (sample In1) increased the RT during the late phase of exhalation, whereas encoding sample cues during the late phase of exhalation (sample Ex2) decreased the RT during the late phase of exhalation.

**Fig. 4.**
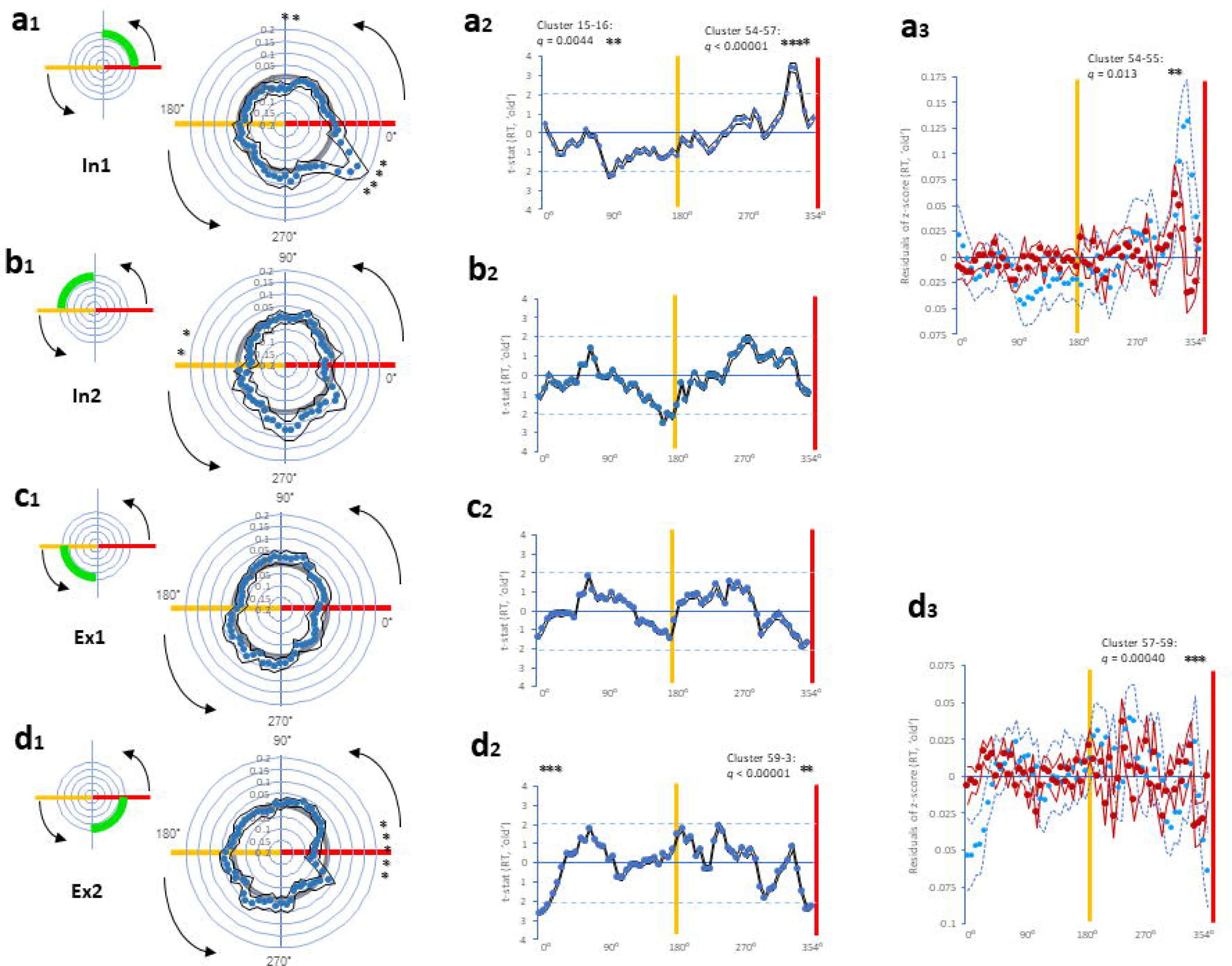
Cluster formations in RT of respiratory phase matching between sample and test blocks (2) Plots showing z-scored RTs (**a_1_**, **b_1_**, **c_1_**, **d_1_**), t-statistics (**a_2_**, **b_2_**, **c_2_**, **d_2_**), and residuals of z-scored RTs (**a_3_**, **d_3_**) of ‘old’ events during the test respiratory cycle, according to four sample respiratory conditions: the sample In1 condition (**a_1_-a_3_**), sample In2 condition (**b_1_, b_2_**), sample Ex1 condition (**c_1_, c_2_**), and sample Ex2 condition (**d_1_-d_3_**). * *p* ≤ 0.05 in **a_1_**, **b_1_**, and **d_1_**. ** *q* ≤ 0.05 (a single cluster with two bins), *** *q* ≤ 0.05 (a single cluster with three bins), **** *q* ≤ 0.05 (a single cluster with four bins) in **a_2_**, **d_2_**, **a_3_**, and **d_3_**.

In addition, we performed residual analysis by dividing the RT datasets of sample events into two conditions (0-180° and 180-360°, Suppl. Fig. S6) and six conditions (0-60°, 60-120°, 120-180°, 180-240°, 240-300°, and 300-360°, Suppl. Fig. S7). ARMA (1, 1) modeling and residual analysis revealed similar results: the sample exhalation condition resulted in cluster formation of residuals during late exhalation in the test block (i.e., the R240/300 condition, Suppl. Fig. S7). These findings confirmed and partially supported the phenomenon of respiratory alignment between encoding and retrieval in the DMTS task.

Moreover, since there were unequal temporal durations between inhalation and exhalation in the circular respiratory analysis, we reduced the number of bins during inhalation by half (i.e., 15 bins) and then conducted the same circular respiratory analysis (Suppl. Fig. S8). The results were obtained as follows: i) residual analysis revealed cluster formation among the RT of test events during late exhalation (Suppl. Fig. S9); ii) encoding sample cues containing the EI transition (inspiratory onset) increased the RT during late exhalation (Suppl. Fig. S10); and iii) the sample condition during early inhalation (sample In1) increased the RT during late exhalation, whereas the sample condition during late exhalation (sample Ex2) decreased the RT during late exhalation (Suppl. Fig. S11). Respiration-timing-dependent analysis of RT with the modified durations between inhalation and exhalation yielded results that were consistent with the main findings.

## Discussion

The present study demonstrated that memory performance was modulated by the phase alignment of respiration between encoding and retrieval in participants performing a DMTS task. Overall, RT was reduced during the late phase of exhalation at retrieval. Moreover, when visual cues were encoded during the late phase of exhalation (sample Ex2), shorter RTs were again observed during late exhalation at retrieval, suggesting a beneficial effect of phase-matched encoding and retrieval during late exhalation. In contrast, longer RTs were observed during late exhalation at retrieval when encoding occurred either i) during the case that encompassed the EI transition (sample EIt) or ii) during the early phase of inhalation (sample In1). These opposing outcomes under identical retrieval conditions indicated that the respiratory phase during encoding is a key determinant of memory performance. These findings underscore the importance of respiratory phase alignment across memory processes, suggesting that optimizing breathing patterns during encoding may shape cognitive efficiency.

We found that encoding during inhalation followed by retrieval during exhalation was linked to diminished cognitive efficiency, whereas both encoding and retrieval during the same exhalation facilitated cognitive performance. These results are consistent with the concept of state-dependent memory, which holds that memory performance is enhanced when the physiological or psychological state at retrieval matches that at encoding[31]. State-dependent memory is the phenomenon whereby information encoded in a particular mental, physiological, psychological, or pharmacological state is more efficiently retrieved when the individual is in a similar state[32,33]. Extending this view, we propose that respiration functions as a phase-dependent internal context that may modulate cognitive load. In particular, the late phase of exhalation may act as an optimal contextual window for both encoding and retrieval, facilitating memory through respiratory phase-locking mechanisms. These results suggest that respiratory phase alignment may play a critical role in shaping the internal contextual framework of cognition.

In this study, we sought to isolate the influence of the respiratory phase during encoding. Growing evidence indicates that respiration modulates memory and emotional processing across distributed brain networks[12,34–37]. However, well-established respiratory effects during retrieval – particularly those linked to inspiratory onset or EI transition[21,22,38] – can obscure respiration-timing-dependent contributions that are specific to encoding. EI-transition-dependent effects exert a disproportionate impact during retrieval, markedly prolonging RTs and degrading accuracy. However, respiratory phase matching between encoding and retrieval has only a modest effect and does not reliably alter accuracy. To minimize the confounding effects of the EI transition during retrieval, we implemented a 2-sec test cycle, which helped prevent interference from the EI transition during retrieval[21]. Consistent with this design, our analysis of RTs aligned to the circular respiratory phase at retrieval revealed clear encoding-dependent modulation of cognitive efficiency across the respiratory cycle.

Our findings showed that cognitive efficiency was comprehensively enhanced at the end of exhalation during retrieval, with a similar improvement emerging around the EI transition. However, our ARMA modeling and residual analysis eliminate the effects of autoregressive and moving average coefficients across adjacent bins, effectively removing history-dependent structures and transient fluctuations in RT and accuracy. Accordingly, the present results do not capture effects that span across the EI transition. Thus, the observed increase in cognitive efficiency was confined to the late phase of exhalation. Consistent with this interpretation, previous studies have reported that successful motor actions are more frequently initiated during late exhalation[16,39]. In particular, late exhalation is characterized by a period of coordinated cortical activity aligned with voluntary motor responses, suggesting that motor actions can be more efficiently executed with reduced cognitive load during late exhalation. However, once the EI transition occurred during encoding, the cognitive load appeared to increase – even during the late exhalation phase of retrieval – emphasizing a potential interaction between the respiratory phase at encoding and subsequent retrieval. Therefore, understanding respiratory conditions during encoding is crucial for predicting cognitive load and modulating memory performance during exhalation.

Moreover, we identified bin clusters of increased and decreased RT and accuracy across the respiratory cycle using ARMA modeling and residual analysis. This approach allows the residuals to reflect white-noise-like fluctuations (i.e., ideal residuals), unmodeled structures (e.g., higher-order dependencies, long-range correlations, and nonlinear dynamics), and rhythmic or periodic components (e.g., respiratory cycle). Since residual fluctuations may still contain rhythmic components, our results suggest that phase-dependent respiratory variations contribute to the temporal dynamics observed in residuals of RT performance.

Several questions arise as to whether respiration-timing-dependent effects reflect temporal coding mechanisms in the brain. Information is conveyed not only by firing rate but also by the precise timing of spikes relative to ongoing neural rhythms or events. Temporal coding is closely associated with neural oscillations, and increasing evidence indicates that respiration can impose structure on these oscillatory dynamics. In rodents, respiratory rhythms entrain neural oscillations across multiple frequency bands – delta (0.5-4 Hz), theta (4-12 Hz), and gamma (30-80 Hz) – via mechanisms as a cross-scale form of cross-frequency coupling, particularly in the olfactory bulb, hippocampus, prefrontal cortex, and widespread cortical areas[40–46]. Although the respiratory frequency varies across species depending on body size (i.e., 3–7 Hz in mice, 1–3 Hz in rats, and 0.15–0.3 Hz in humans during wakefulness), respiration-locked activity is also observed in the human brain. For example, intracranial EEG (iEEG) recordings in epilepsy patients have revealed respiration-coupled oscillations in the piriform cortex and hippocampus[12,17]. In a large-scale analysis, a form of interoceptive-neural oscillatory coupling between respiration (0.15-0.3 Hz) and hippocampal local field potentials was identified, peaking at 0.28 Hz[47]. These findings suggest that the slower respiratory rhythms in humans may organize perception, cognition, and behavior by modulating the phase-specific timing of neuronal firing.

These findings may extend beyond desk-based cognitive tasks to outdoor physical activity and fieldwork, where specific breathing strategies could similarly be applied. During training or encoding, the optimal timing for learning information appears to occur during late exhalation. However, once respiration proceeds beyond inspiratory onset (EI transition), this phase may become suboptimal for task performance, as the cognitive load tends to increase. Specifically, retrieval encompassing inspiratory onset (EI transition) is strongly associated with both reduced accuracy and prolonged RTs, particularly when the test cycle duration exceeds 3 sec, allowing phase-locking to the respiratory rhythm[21,23]. Therefore, training from the outset in a manner that avoids reaching the inspiratory onset (EI transition) during both encoding and retrieval is essential. This can be achieved by learning to prolong exhalation as much as possible. Indeed, extending exhalation during both encoding and retrieval significantly reduced the occurrence of inspiratory onset (EI transition) and increased the proportion of test trials that remained fully with the exhalation phase (data not shown). By systematically training to prolong exhalation, individuals may expand the temporal window for effective encoding and enhance resilience to inspiratory-onset interference during both learning (encoding) and testing (retrieval).

## Methods

### Subjects

The participants in this study were 31 healthy volunteers. None of the participants were regularly taking medication, and none had a known history of respiratory, cardiovascular, endocrine, neurological, or psychiatric disease. Written informed consent was obtained from all participants. All participants were scored as right-handed according to the Flanders Handedness Survey[48]. One subject was excluded because it did not reach the criterion (the hit score was less than 60%). In total, 30 healthy participants (18 females and 12 males; age: 22.17 ± 0.37 years, range: 20-27 years) were used for further analyses. All procedures performed on humans were in accordance with the Declaration of Helsinki (Ethical Principles for Medical Research Involving Hunan Subjects) and the Ethical Guidelines for Medical and Health Research Involving Human Subjects, Japan, and all procedures were approved by the Ethics Committee of Hyogo Medical University, Japan (1825).

### Physiological apparatus

Inhalation and exhalation during the respiratory cycle were continuously recorded via a flow sensor nasal cannula (Flow Nasal Cannula A, Atom Medical, Japan) equipped with a differential pressure transmitter (Model KL17, Nagano Keiki, Japan). Electrocardiograms were recorded via lead II (voltage between the left leg and right arm electrodes) using a differential biological amplifier (Bioamp, AD Instruments, New Zealand). These physiological parameters and cognitive parameters including temporal information of visual cues and button-press responses for the task were sampled at 1 kHz via the PowerLab data acquisition system (PowerLab, AD Instruments) and were processed online via LabChart software (LabChart 8.1, AD Instruments). Before each experiment, the air pressure in the experimental room was measured to determine the baseline level.

### Behavioral task paradigm

A DMTS version of a visual recognition memory task with a short delay was employed as previously described by Nakamura et al.[21,22] with modifications. The DMTS task consists of a sample block, delay block, and test block structured according to a standard DMTS protocol[1]. Task paradigms were created in NBS Presentation software (Presentation®, Neurobehavioral Systems). The DMTS task required the memorization and recognition of natural objects (or items) in the photographs, which were randomly selected from 26,107 photographs in the THINGS database[29,30]. In each session, individual participants memorized 40 photographs (sample cues), discriminated 80 photographs (test cues, Fig. 1A), and repeated the session 10 times (in total, 400 memorizations and 800 discriminations per participant). The volunteers were instructed to fix their eyes on the white cross at the center of the screen. At the beginning of the sample block, the white cross turned green, and each participant was then exposed to a series of 40 photographs (i.e., 40 sample cues) displayed one at a time on the screen. Each sample cue was continuously presented on the screen every 1 sec and each sample event lasted 1 sec (Fig. 1B).

After a delay (20-21 s), the white cross turned red, and then each participant was tested 80 separate times for the ability to distinguish between visual cues presented during the sample block (‘old’ or match cues) and ‘new’ (or nonmatch) cues. After each test cue was presented, the volunteers pressed one of two buttons using their fingers once they had identified the cue shown during the test block. 40 ‘old’ cues and 40 ‘new’ cues were presented in a random manner during the test block. The interstimulus interval of the test cues was approximately 2 sec, with one of three different time lags in random order (0, 150, 300 ms after a trigger signal). Before the experiments, the volunteers were instructed on how to perform the task and were told to breathe in a relaxed, natural manner during the task. We confirmed that the volunteers did not intentionally control their breathing during the task, since no volunteers reported awareness of the cues being locked to their own respiratory cycle or of intentionally adjusting their breathing to correspond with the timing of cue presentation to improve their performance.

### Data processing

During the experiments, we measured 1) cognitive parameters including the type and timing of cue presentation and button presses, via NBS Presentation software; and 2) respiratory parameters, including the onset of inhalation and exhalation, via LabChart software.

A test event and its RT were defined as the period from test cue presentation to a button-press response. Test events were classified into the following conditions (Fig. 2B): (i) test events that occurred entirely within inhalation without the transitions (the test NoI or NoI_t_ condition); (ii) test events that encompassed the inhalation-to-exhalation (IE) transition (the test IEt or IEt_t_ condition); (iii) test events that occurred entirely within exhalation without the transitions (the test NoE or NoE_t_ condition); and (iv) test events that encompassed the exhalation-to-inhalation (EI) transition (the test EIt or EIt_t_ condition). Notably, test events that included both the IE and EI transitions (a condition designated ‘double’) were excluded from further analysis because of an insufficient number of occurrences (Fig. 2B).

A sample event was defined as a 1-sec period corresponding to each sample cue presentation. Sample events were divided into the following conditions (Fig. 2G): (i) sample events that occurred within inhalation without the transitions (the sample NoI or NoI_s_ condition); (ii) sample events that encompassed the IE transition (the sample IEt or IEt_s_ condition); (iii) sample events that occurred within exhalation without the transitions (the sample NoE or NoE_s_ condition); and (iv) sample events that encompassed the EI transition (the sample EIt or EIt_s_ condition). On the basis of the classification of sample events, test events were further subdivided into the following conditions: the NoI_s_ series: (i) NoI_s_-NoI_t_, (ii) NoI_s_-IEt_t_, (iii) NoI_s_-NoE_t_, and (iv) NoI_s_-EIt_t_ conditions; the IEt_s_ series: (v) IEt_s_-NoI_t_, (vi) IEt_s_-IEt_t_, (vii) IEt_s_-NoE_t_, and (viii) IEt_s_-EIt_t_ conditions; the NoE_s_ series: (ix) NoE_s_-NoI_t_, (x) NoE_s_-IEt_t_, (xi) NoE_s_-NoE_t_, and (xii) NoE_s_-EIt_t_ conditions; and the EIt_s_ series: (xiii) EIt_s_-NoI_t_, (ivx) EIt_s_-IEt_t_, (vx) EIt_s_-NoE_t_, and (vix) EIt_s_-EIt_t_ conditions. We excluded sample and/or test events that encompassed both the IE and EI transitions (a condition designated ‘double’, Fig. 2H). Sample events were also categorized as follows: (i) the sample In1 (overlapping in the 0-90° bin), (ii) sample In2 (overlapping in the 90-180° bin), (iii) sample Ex1 (overlapping in the 180-270° bin), and (iv) sample Ex2 (overlapping in the 270-360° bin) conditions.

Regarding the circular respiratory bin-dependent analysis, datasets of RT and accuracy were z-scored within each respiratory cycle for each participant. To determine significant phase-locked patterns of RT and accuracy during the respiratory cycle, each value was assigned to a corresponding respiratory phase bin. The circular respiratory cycle, which was defined as 0-180° for inhalation and 180-360° for exhalation, was divided into 60 nonoverlapping bins (6° per bin, from 0° to 360°). Because test events or RTs span time, a single event could contribute to multiple adjacent bins because of temporal overlap. Importantly, although the Hilbert transform is useful for estimating continuous and instantaneous phases, it does not ensure consistent alignment with physiologically defined events, such as inhalation onset (EI transition) or exhalation onset (IE transitions). Furthermore, as the distributions of sample events, test events, and ‘old’ events formed smooth circular patterns (Fig. 1 G-I), we did not apply the Hilbert transform in the present study.

### Statistical analysis

All the statistical analyses were performed using R version 4.5 software (R Foundation for Statistical Computing, 2025; https://www.r-project.org/). Across individual participants, RT and accuracy were tested for normality and sphericity via the Shapiro-Wilk test for normality and the Mauchly test for sphericity. We used one-way repeated-measures ANOVA, and post hoc pairwise comparisons via paired *t* tests with the Bonferroni correction. If the assumption of normality was violated, we carried out the nonparametric Friedman test followed by post hoc comparisons via the Wilcoxon signed-rank test with the Bonferroni correction. Circular respiratory bin-dependent analysis is described below.

### Circular bin-dependent analysis step 1

For circular respiratory bin-dependent analysis for z-scored RT and z-scored accuracy, a two-tailed one-sample *t* test against zero was performed for each bin (n = 30 participants). Two or more adjacent bins, each exceeding the significance threshold (*p* ≤ 0.05, uncorrected), were defined as a single candidate cluster. For each cluster, the sum of the *t* values was used for cluster-level analysis: a nonparametric cluster-based permutation test (100,000 iterations) was conducted by randomly flipping the sign of each participant’s 60-bin vector, thereby generating a null distribution of cluster-level *t*-sums. FDR correction was applied across all clusters to control for multiple comparisons.

### Circular bin-dependent analysis step 2

Since datasets of RT and accuracy across adjacent phase bins are not statistically independent due to the temporal smoothing imposed by overlapping their durations, we employed an ARMA model. For each participant, a time series of z-scored RT or z-scored accuracy in 60 bins was modeled via the autoregressive integrated moving average (ARIMA) function (*p* = 1, *d* = 0, *q* = 1, forecast package in R), and the autocorrelated moving structure was subsequently removed from the datasets. To validate our choice of the ARMA (*p* = 1, *q* = 1) model for detrending the binwise series of RT or accuracy, we compared its performance to that of the model selected by the auto.arima() function based on the Akaike information criterion (AIC) and Bayesian information criterion (BIC). The ARMA (1, 1) model demonstrated comparable or superior fit, justifying its use in the subsequent residual analysis. The resulting residuals were again estimated under the same cluster-based statistical procedure (two-tailed one-sample *t* test per bin, cluster formation, permutation test, and FDR correction). For the normality assumption required for *t* tests, Shapiro-Wilk tests and Q-Q plots were used. While the Shapiro-Wilk test rarely yielded significant results (n = 30 participants), the Q-Q plots demonstrated that the residuals were approximately normally distributed, particularly in the central range. Given the *t* test’s known robustness to mild deviations from normality and the visual confirmation, we proceeded with parametric testing.

## Acknowledgments

We thank Norihiro Sadato (National Institute for Physiological Sciences, Okazaki, Japan), Yoshitaka Oku (Hyogo Medical University, Nishinomiya, Japan) for comments on and discussion of the manuscript, and Toshiro Saito (Hyogo Medical University) for experimental supports.

## Author contributions

NHN and MF contributed to the study design; NHN contributed to acquisition of data; NHN, KY, and MF contributed to methodology; NHN and KY contributed to data analysis; NHN and KY contributed to data interpretation; NHN contributed to drafting of manuscript; NHN, KY and MF contributed to the final manuscript.

## Data availability

The data that support the main findings of this study are available from the corresponding author upon request. The datasets used in this study are also available at https://github.com/nakamunh/Nakamura_2025_Sci_Rep.

