## Supplemental Figures for "Breath as an internal context: Respiratory phase alignment between encoding and retrieval optimizes memory performance"

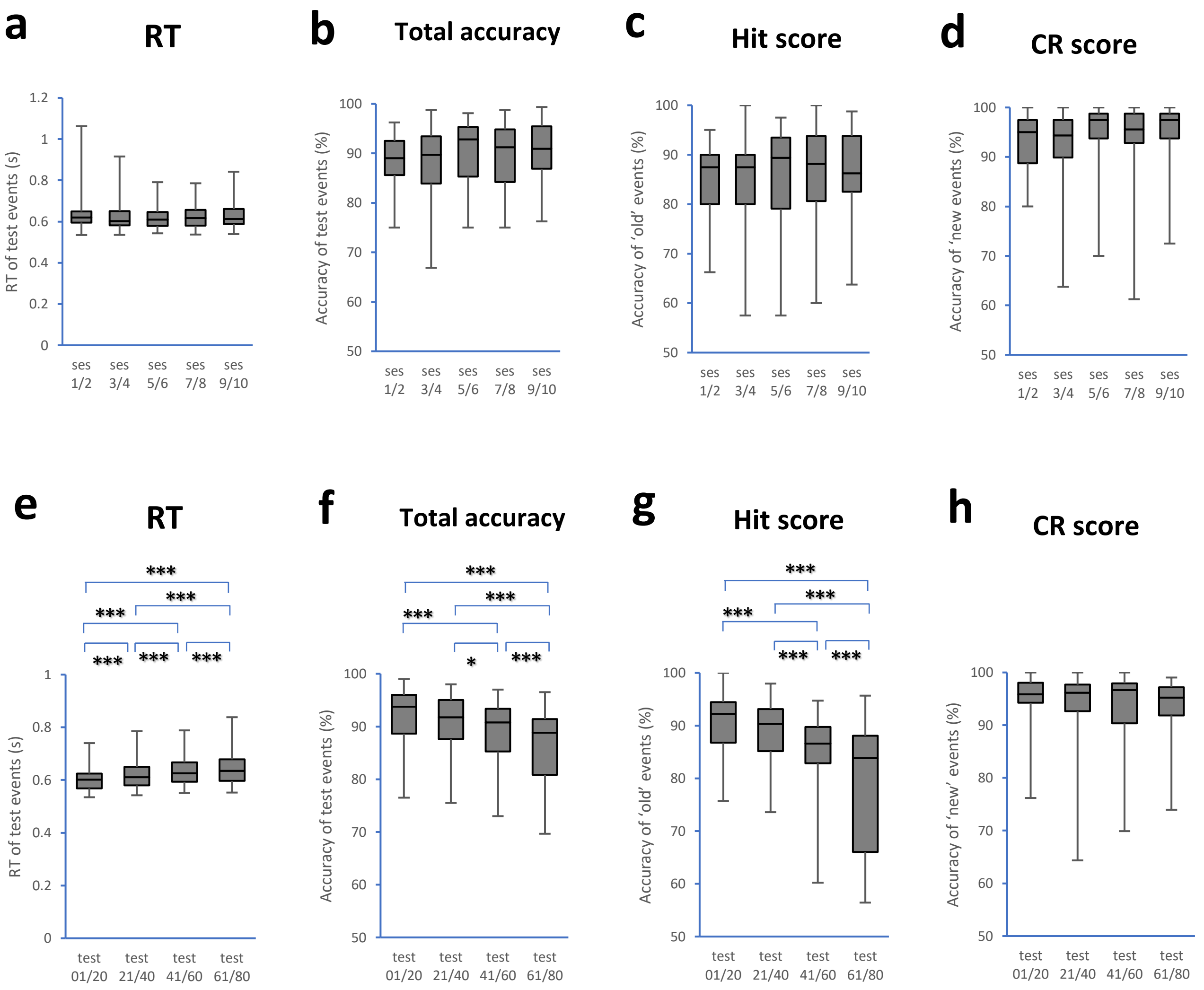


**Fig. S1 Task performance among sessions and within a session**

Individual participants memorized 40 photographs and discriminated 80 photographs in each session, and then repeated the session 10 times. To determine the differences of task performance among sessions, test events were classified into the following conditions: (i) sessions 1 and 2 (Ses 1/2); (ii) sessions 3 and 4(Ses 3/4); (iii) sessions 5 and 6 (Ses 5/6); (iv) sessions 7 and 8 (Ses 7/8); and (v) sessions 9 and 10 (ses 9/10).There were no differences in RT of test events ($\chi^2(4) = 3.95$, $p = 0.41$, Friedman test, **a**), total accuracy (accuracy of test events, $\chi^2(4) = 9.25$, $p = 0.0552$, **b**), Hit score (accuracy of 'old' events, $\chi^2(4) = 5.48$, $p = 0.24$, **c**), while correct rejection (CR) score (accuracy of 'new' events) had the significant difference ($\chi^2(4) = 13.34$, $p = 0.0097$, **d**). However, post hoc comparisons did not show any difference of CR score among the conditions (Wilcoxon signed rank test with Bonferroni corrections).

To identify the differences of task performance within a session, test events were classified into (i) test events 01 to 20 (Test 01/20); (ii) test events 21 to 40 (Test 21/40); (iii) test events 41 to 60 (Test 41/60); (iv) test events 61 to 80 (Test 61/80). The Shapiro-Wilk normality test did not show normal distributions of RT and accuracy of test events. There were significant differences in RT of test events ($\chi^2(3) = 68.32$, $p = 9.8 \times 10^{-15}$, Friedman test, **e**), total accuracy of test events ($\chi^2(3) = 60.92$, $p = 3.8 \times 10^{-13}$, **f**), Hit score ($\chi^2(3) = 66.08$, $p = 3.0 \times 10^{-14}$, **g**), and CR score ($\chi^2(3) = 8.12$, $p = 0.044$, **h**). Post hoc comparisons showed that the Test 01/20 condition exhibited in the lowest RT, the highest accuracy, and the highest Hit score, whereas the Test 61/80 condition had the highest RT, the lowest accuracy, and the lowest Hit score. No difference of CR score was observed. * $p \le 0.05$; ** $p \le 0.01$; and *** $p \le 0.005$ (Wilcoxon signed rank test with Bonferroni corrections).

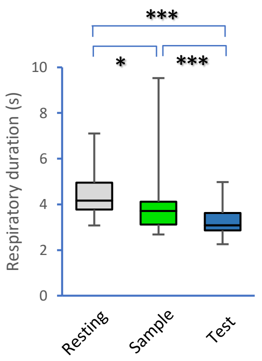


**Fig. S2 Respiratory dynamics during the task**

Plots showing durations of respiration among the resting state, the sample block, and test block ($\chi^2(2) = 41.267$, $p = 1.1 \times 10^{-9}$, Friedman test). Post hoc comparisons revealed that the test block presented the shortest durations of respiration, and the sample block presented the second shortest durations of respiration (Wilcoxon signed rank test with Bonferroni correction).

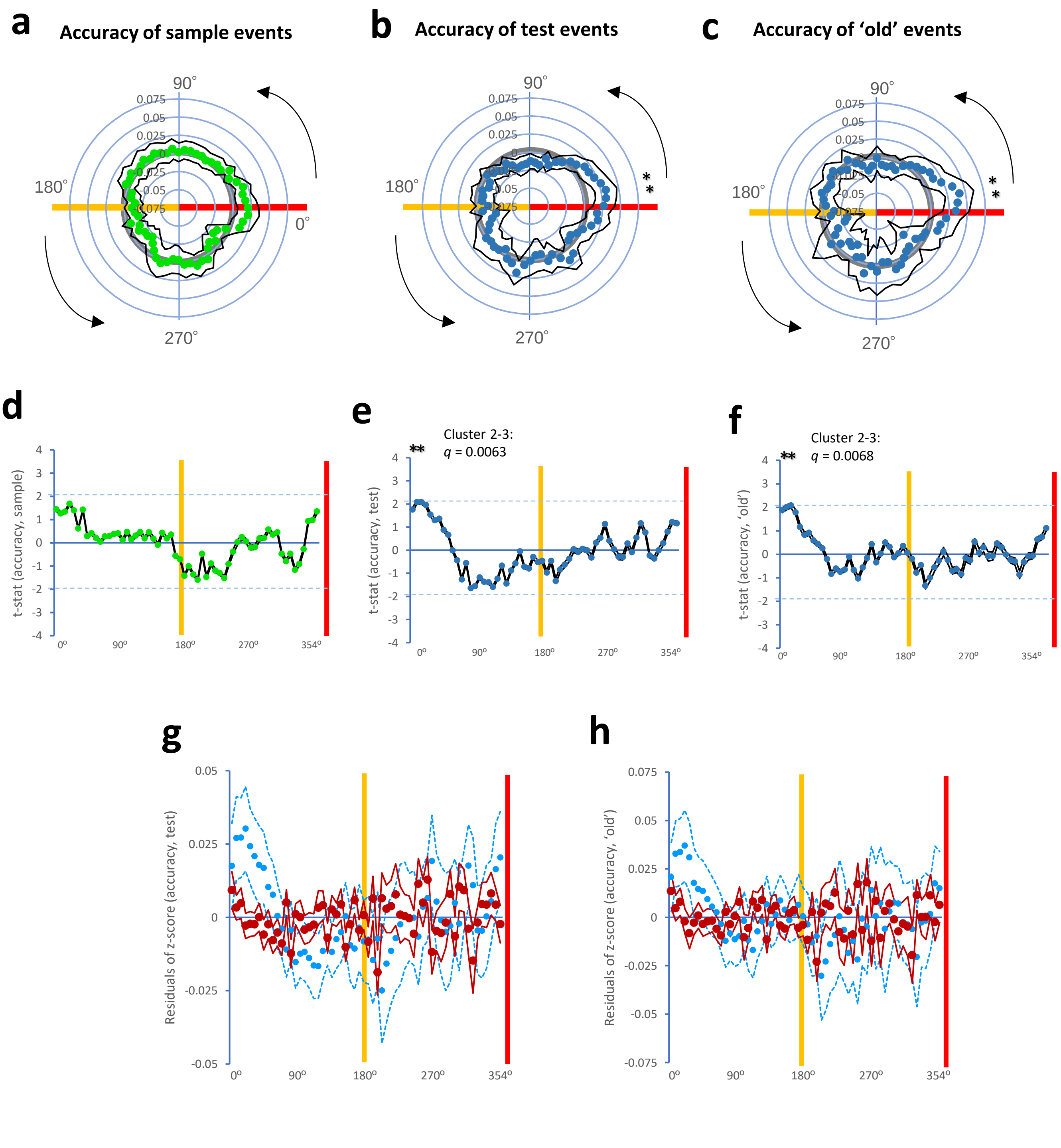


**Fig. S3  Respiration-timing-dependent accuracy variations**

To investigate cluster formations in accuracy along the respiratory cycles, we used respiratory phase (bin)-dependent analysis in accuracy of sample events, test events, and 'old' events. No skewed distribution of z-scored accuracy of sample events was observed along the sample respiratory cycle (green dots, **a**), There were skewed distributions of z-scored accuracy of test events along the test respiratory cycle (dark blue dots, *ps* $\leq$ 0.05, two-tailed one sample *t*-test against zero, **b**), and accuracy of 'old' events along the test respiratory cycle (dark blue dots, *ps* $\leq$ 0.05, **c**). Nonparametric permutation test and FDR correction showed the significance of cluster formations in accuracy of test events (cluster 2-3: $t_{(sum)}$ = 4.15, $q$ = 0.0063, **e**), and accuracy of 'old' events (cluster 2-3: $t_{(sum)}$ = 4.08, $q$ = 0.0068, **f**). Meanwhile, no difference in accuracy of sample events was observed (**d**). ARMA ($p$ = 1, $q$ = 1) modeling and residual analysis using permutation test and FDR correction did not show cluster formations of residuals (dark red dots) in accuracy of test events (**g**), nor accuracy of 'old' events (**h**). Light blue dots in **g** and **h** indicate ARMA (1, 1) fitted dots. * $p \leq 0.05$ (Two-tailed one-sample *t*-test against zero) in **a-c**, ** $q \leq 0.05$ (a single cluster with two bins, permutation test and FDR correction) in **d-h**.

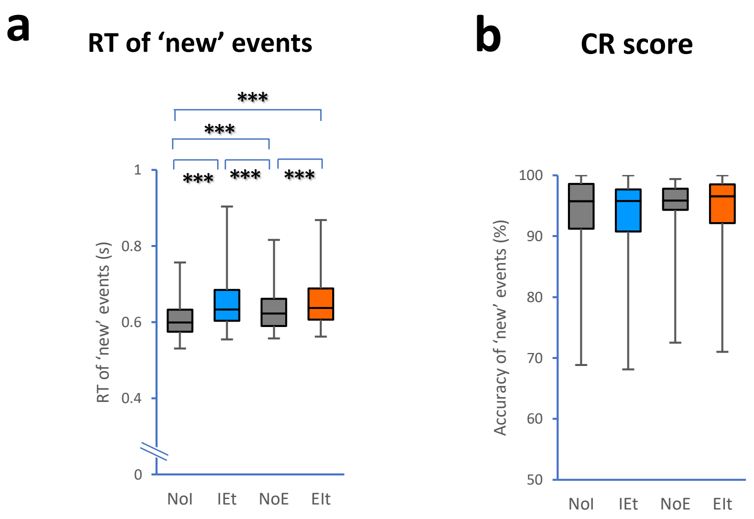


**Fig. S4  RT of 'new' events and CR score among test block-dependent respiratory conditions**

Test events were classified into (i) the NoI condition; (ii) the IEt condition; (iii) the NoE condition; and (iv) the EIt condition along the test respiratory cycle. The Shapiro-Wilk normality test did not show normal distributions of  RT of 'new' events and correct rejection (CR) score (accuracy of 'new' events). There were significant differences in RT of 'new' events ($\chi^2(3) = 63.44$, $p = 1.1 \times 10^{-13}$, Friedman test, **a**). However, no difference in CR score (accuracy of 'new' events) was observed ($\chi^2(3) = 4.04$, $p = 0.26$, **b**). Post hoc comparisons showed that both the IEt and EIt conditions exhibited in the highest RT of 'new' events, whereas NoI condition had the lowest RT of 'new' events (Wilcoxon signed rank test with Bonferroni corrections). *** $p \leq 0.005$ (Wilcoxon signed rank test with Bonferroni corrections).

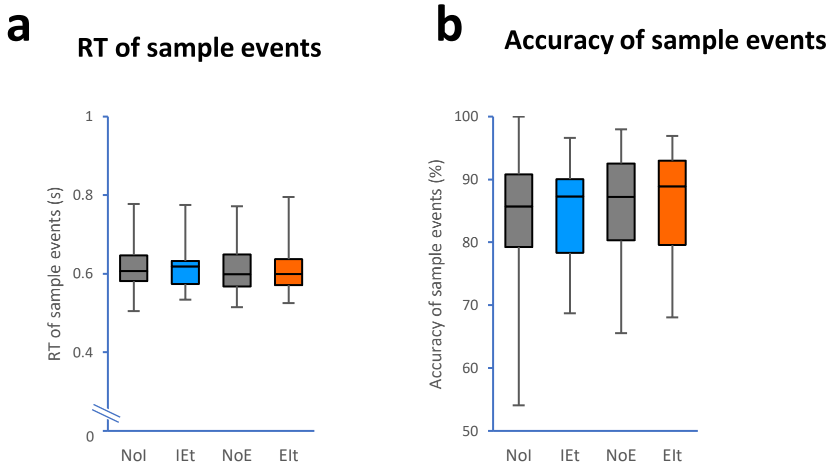


**Fig. S5 RT and accuracy of sample events among sample block-dependent respiratory conditions**

Sample events were classified into (i) the NoI condition; (ii) the IEt condition; (iii) the NoE condition; and (iv) the EIt condition along the sample respiratory cycle. The Shapiro-Wilk normality test did not show normal distributions of RT nor accuracy of sample events. No differences in RT of sample events ($\chi^2(3) = 3.04$, $p = 0.39$, Friedman test, **a**) and accuracy of sample events ($\chi^2(3) = 2.16$, $p = 0.54$, **b**) were observed among the sample respiratory conditions.

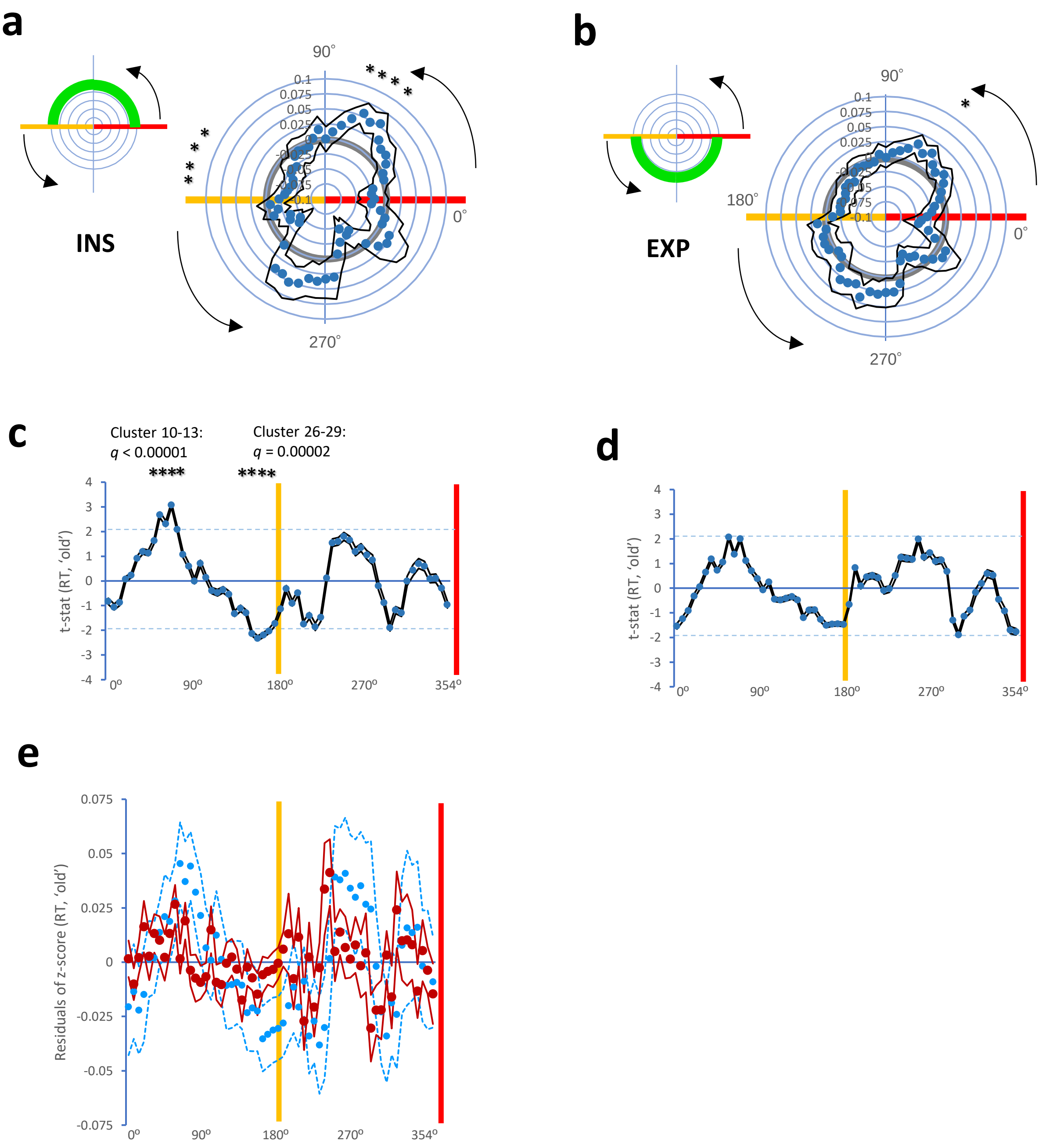


**Fig. S6 Cluster formations in RT of respiratory phase matching between sample and test blocks (3)**

We divided the RT dataset of sample events into: (i) sample INS (the overlap of sample events in 0-180º bin); and sample EXP (the overlap of sample events in 180-360º bin) conditions. We found skewed distribution of z-scored RT of sample INS (**a**) and EXP (**b**) conditions along the test respiratory cycle (dark blue dots, *ps* ≤ 0.05, two-tailed one sample *t*-test against zero). Nonparametric permutation test and FDR correction showed the significance of cluster formations in RT of sample INS condition (cluster 10-13: $t_{(sum)}$ = 10.16, $q < 1.0 \times 10^{-5}$; cluster 26-28: $t_{(sum)}$ = 8.66, $q = 2.0 \times 10^{-5}$, **c**). No cluster formation in RT of sample EXP condition was observed (**d**). However, ARMA (1, 1) modeling and residual did not show cluster formations of residuals (dark red dots) in RT of sample INS condition (**e**). Light blue dots in **e** indicate ARMA (1, 1) fitted dots. * $p \leq 0.05$ (Two-tailed one-sample *t*-test against zero), *** $q \leq 0.05$ (a single cluster of three bins, permutation test and FDR correction); **** $q \leq 0.05$ (each bin for a single cluster).

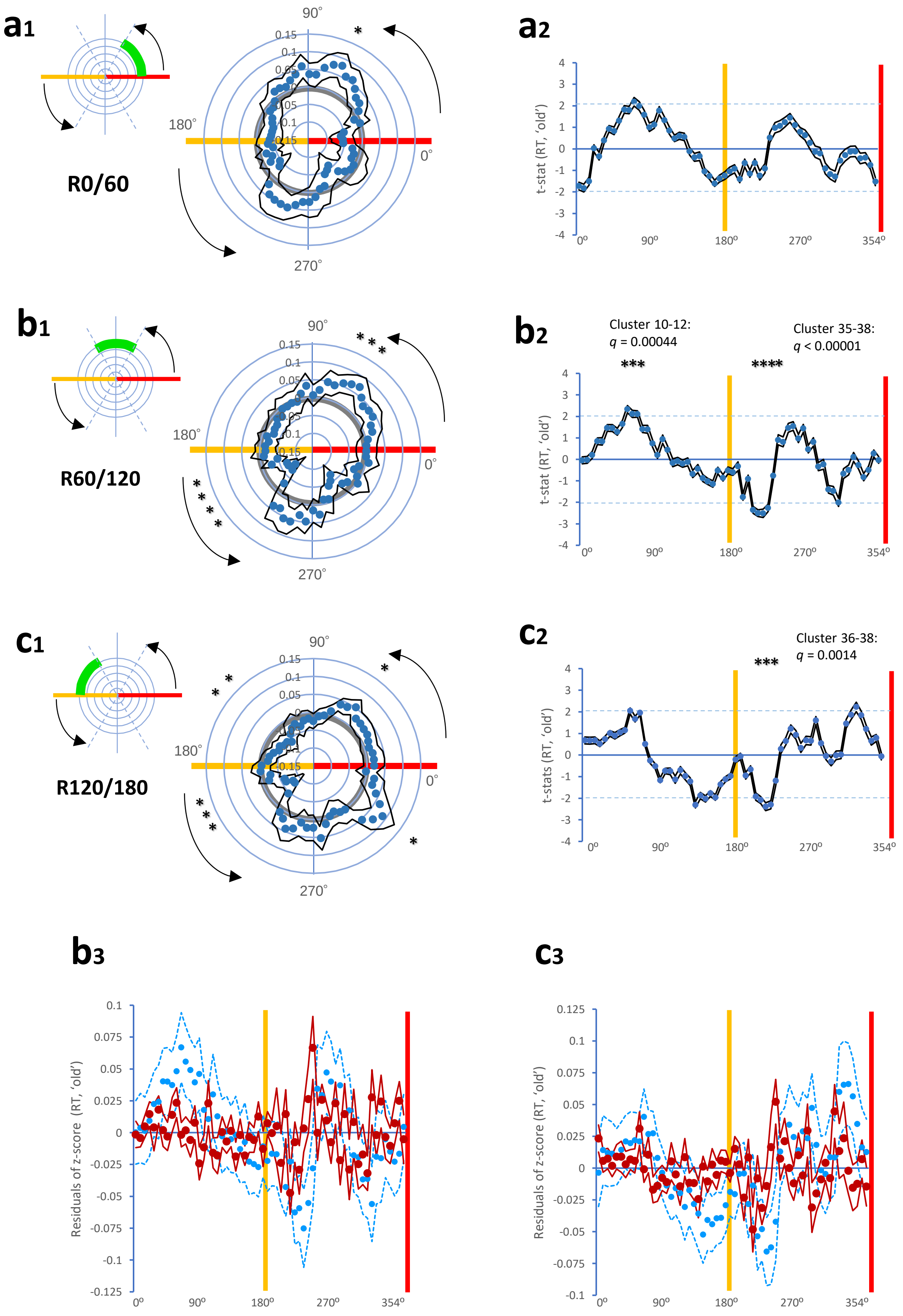
a1
R0/60
90°
180°
0°
270°
0.15
0.1
0.05
*
a2
t-stat (RT, 'old')
4
3
2
1
0
-1
-2
-3
-4
0°
90°
180°
270°
354°
b1
R60/120
*
*
*
*
*
*
*
b2
Cluster 10-12:
q = 0.00044
Cluster 35-38:
q < 0.00001
***
****
t-stat (RT, 'old')
c1
R120/180
*
*
*
*
*
*
*
*
c2
Cluster 36-38:
q = 0.0014
***
t-stats (RT, 'old')
b3
Residuals of z-score (RT, 'old')
0.1
0.075
0.05
0.025
0
-0.025
-0.05
-0.075
-0.1
-0.125
c3
0.125
0.1
0.075
0.05
0.025
0
-0.025
-0.05
-0.075
-0.1

d1

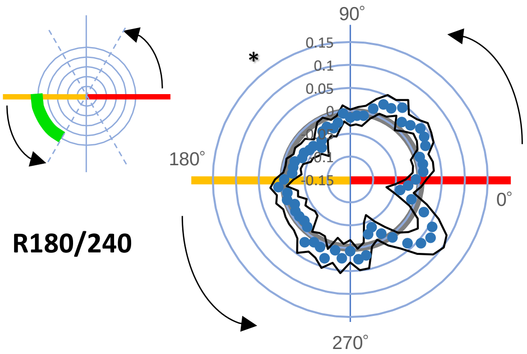


d2

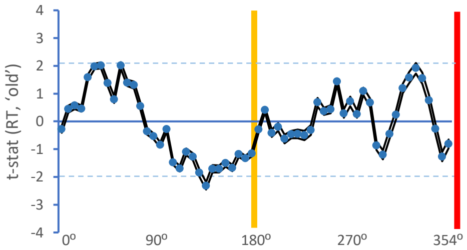


e1

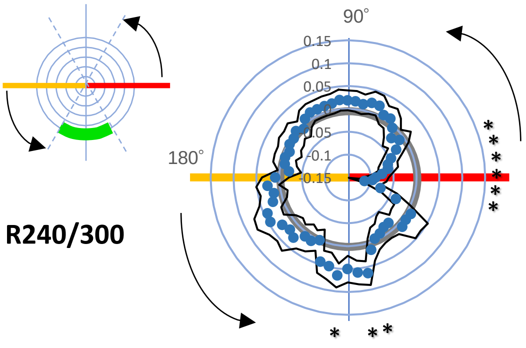


e2

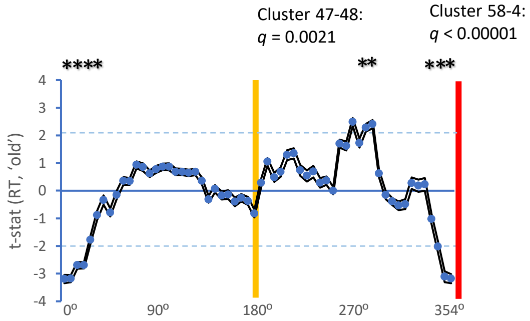


f1

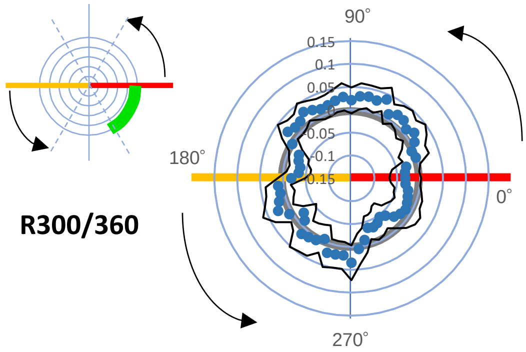


f2

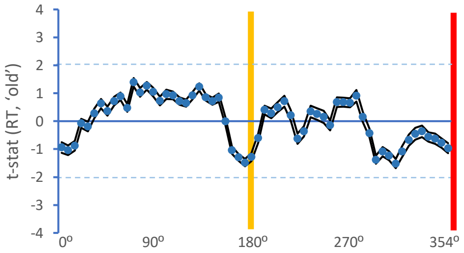


e3

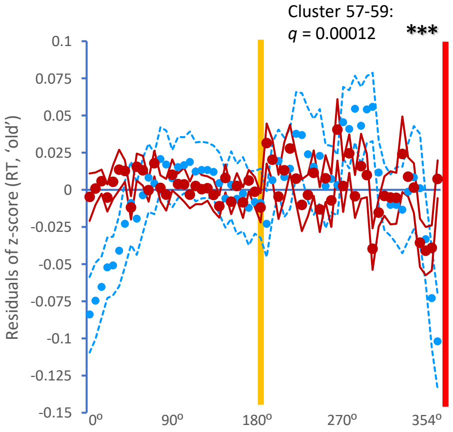

**Fig. S7 Cluster formations in RT of respiratory phase matching between sample and test blocks (4)**

We divided the RT dataset of sample events into: (i) sample R0/60 (the overlap of sample events in 0-60º bin, **a**); (ii) sample R60/120 (the overlap of sample events in 60-120º bin, **b**); (iii) sample R120/180 (the overlap of sample events in 120-180º bin, **c**); (iv) sample R180/240 (the overlap of sample events in 180-240º bin, **d**); (v) sample R240/300 (the overlap of sample events in 240-300º bin, **e**); and (vi) sample R300/360 (the overlap of sample events in 300-360º bin, **f**) conditions. We found skewed distribution of z-scored RT of these conditions along the test respiratory cycle (dark blue dots, *ps* ≤ 0.05, two-tailed one sample *t*-test against zero). Nonparametric permutation test and FDR correction showed the significance of cluster formations in RT of sample R60/120 condition (cluster 10-12: $t_{(sum)} = 6.56$, $q = 0.00044$; cluster 35-38: $t_{(sum)} = 9.63$, $q < 1.0 \times 10^{-5}$, $\mathbf{b_2}$), sample R120/180 condition (cluster 36-38: $t_{(sum)} = 6.78$, $q = 0.0014$, $\mathbf{c_2}$), and sample R240/300 condition (cluster 47-48: $t_{(sum)} = 4.68$, $q = 0.0021$; cluster 58-4: $t_{(sum)} = 20.08$, $q < 1.0 \times 10^{-5}$, $\mathbf{e_2}$). ARMA (1, 1) modeling and residual analysis showed cluster formations of residuals (dark red dots) in RT of sample R240/300 condition (cluster 57-59: $t_{(sum)} = 7.31$, $q = 0.00012$, $\mathbf{d_3}$), but not in RT of sample R60/120 condition ($\mathbf{b_3}$) nor sample R120/180 condition ($\mathbf{c_3}$). Light blue dots in $\mathbf{b_3}$, $\mathbf{c_3}$ and $\mathbf{e_3}$ indicate ARMA (1, 1) fitted dots. * $p \leq 0.05$ (Two-tailed one-sample *t*-test against zero), ** $q \leq 0.05$ (a single cluster of two bins, permutation test and FDR correction); *** $q \leq 0.05$ (each bin for a single cluster); and **** $q \leq 0.05$ (a single cluster of four bins).

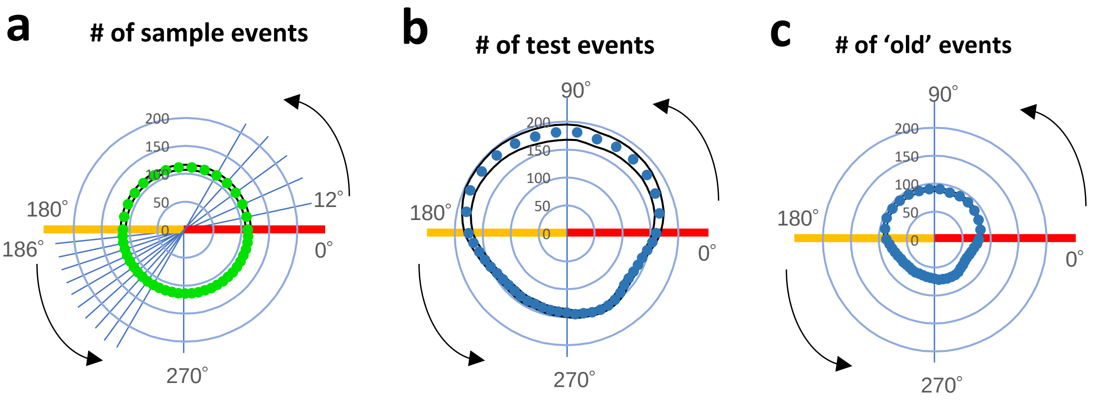


**Fig. S8. Re-binned circular distribution of the respiratory phase**

**a-c**. The inhalation phase was re-binned at half the original resolution (i.e., each 12º bin) to account for the unequal temporal durations between inhalation and exhalation. Plots showing circular histograms of sample events (green, **a**), test events (dark blue, **b**), and 'old' events (dark blue, **c**), where the overlap of individual events with each bin was counted throughout the respiratory cycle. The cognitive data were plotted in a total of 45 bins (15 bins during inhalation: 0-180º; 30 bins during exhalation: 180-360º). The yellow and red lines in **a-c** indicate the IE transition (exhalation onset) and EI transition (inhalation onset), respectively.

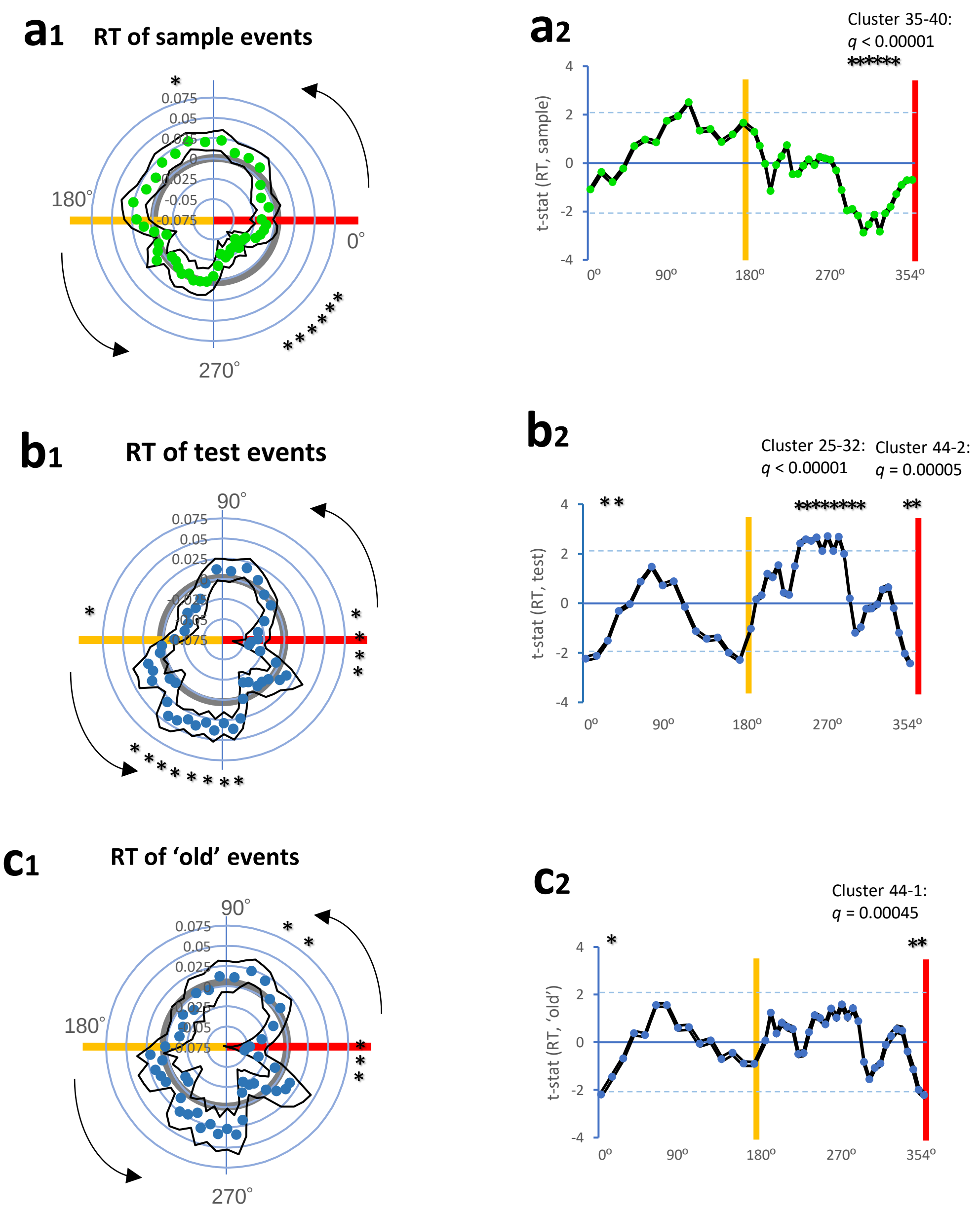
a1
RT of sample events
180°
0°
270°
a2
Cluster 35-40:
q < 0.00001
t-stat (RT, sample)
0°
90°
180°
270°
354°
b1
RT of test events
90°
b2
Cluster 25-32:
q < 0.00001
Cluster 44-2:
q = 0.00005
t-stat (RT, test)
0°
90°
180°
270°
354°
c1
RT of 'old' events
90°
180°
270°
c2
Cluster 44-1:
q = 0.00045
t-stat (RT, 'old')
0°
90°
180°
270°
354°

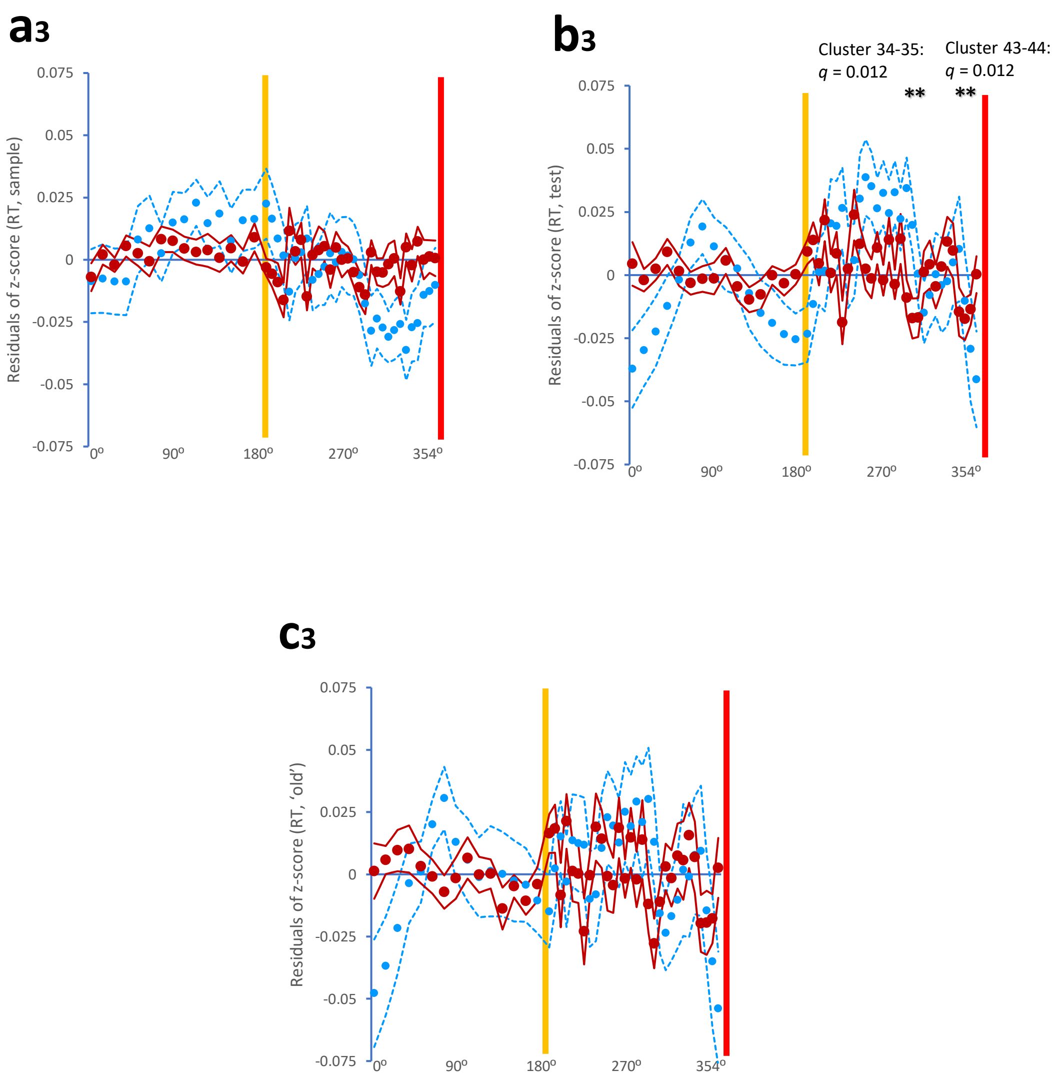


**Fig. S9. Re-binned analysis of respiration-timing-dependent RT variations**

**$a_1$-$c_1$**. Plots showing re-binned circular distributions of z-scored RTs of sample events (**$a_1$**), test events (**$b_1$**), and 'old' events (**$c_1$**) in the respiratory cycle. We found skewed distributions of z-scored RT of sample events, test events, and 'old' events (*ps* ≤ 0.05, two-tailed one sample *t*-test against zero). **$a_2$-$c_2$**. Plots showing t-stats of z-scored RTs of sample events (**$a_2$**), test events (**$b_2$**) and 'old' events (**$c_2$**) together with cluster formations. The nonparametric permutation test and FDR correction revealed the significance of cluster formation (more than two adjacent bins) in the RT of sample events (cluster 35-40: $t_{(sum)}$ = 14.65, $q < 1.0 \times 10^{-5}$), test events (cluster 25-32: $t_{(sum)}$ = 19.80, $q < 1.0 \times 10^{-5}$; cluster 44-2: $t_{(sum)}$ = 8.84, $q = 4.5 \times 10^{-5}$), and 'old' events (cluster 44-1: $t_{(sum)}$ = 6.41, $q$ = 0.00045). **$a_3$-$c_3$**. Plots showing residuals (dark red) of z-scored RTs of sample events (**$a_3$**), test events (**$b_3$**), and 'old' events (**$c_3$**) together with cluster formations. We found that residuals formed clusters in RT of test events (cluster 34-35: $t_{(sum)}$ = 4.50, $q$ = 0.012; cluster 43-44: $t_{(sum)}$ = 4.47, $q$ = 0.012). The light blue dots indicate the ARMA (1, 1) fitted dots. Yellow and red lines in **a-c** indicate the IE transition (exhalation onset) and EI transition (inhalation onset), respectively. * $p \leq 0.05$ in **$a_1$-$c_1$**, ** $q \leq 0.05$ (a single cluster with two bins), *** $q \leq 0.05$ (a single cluster with three bins), and **** $q \leq 0.05$ (a single cluster with four bins).

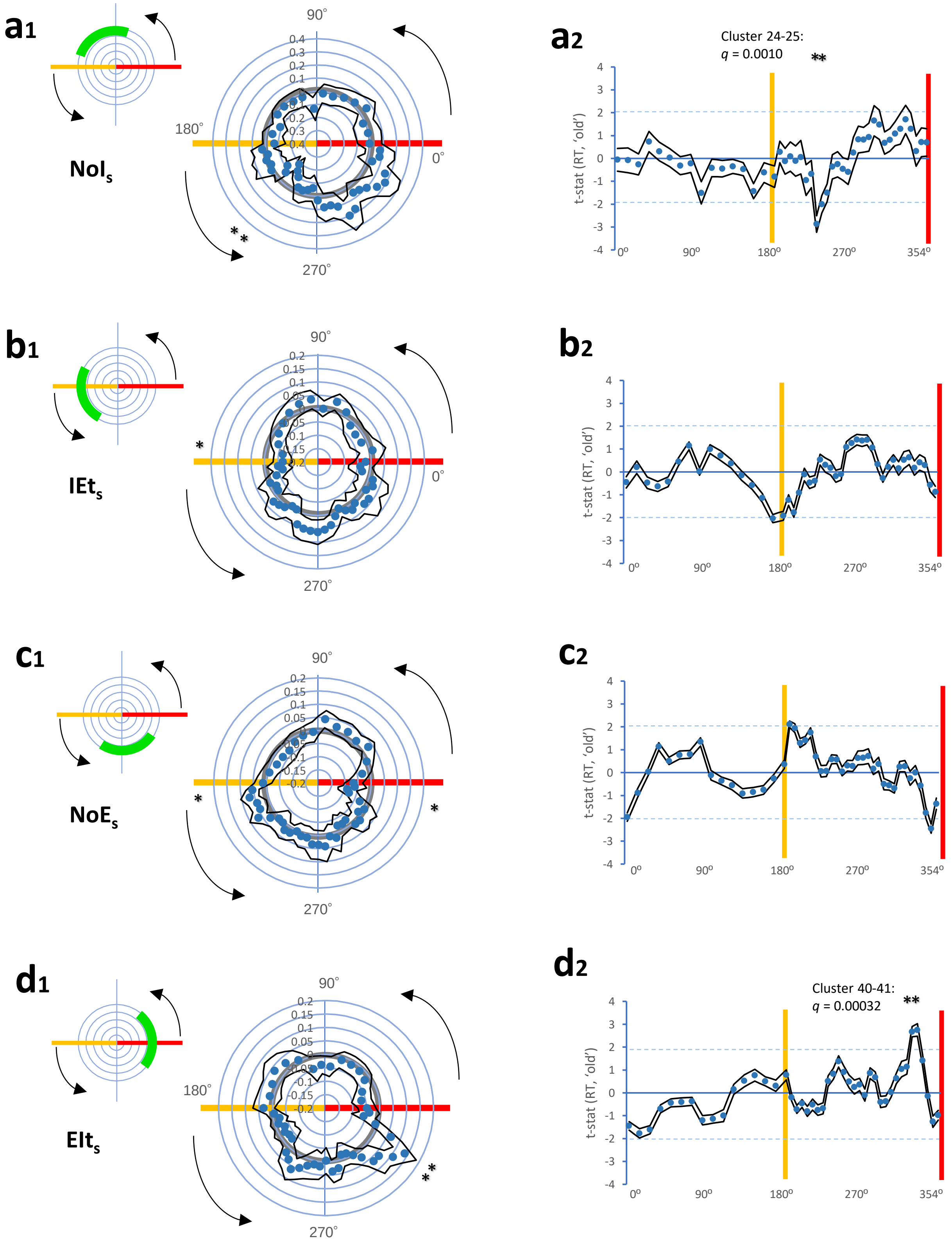

a1
NoIs
a2
Cluster 24-25:
q = 0.0010
**
t-stat (RT, 'old')
b1
IEts
b2
c1
NoEs
c2
d1
EIts
d2
Cluster 40-41:
q = 0.00032
90°
180°
0°
270°
354°

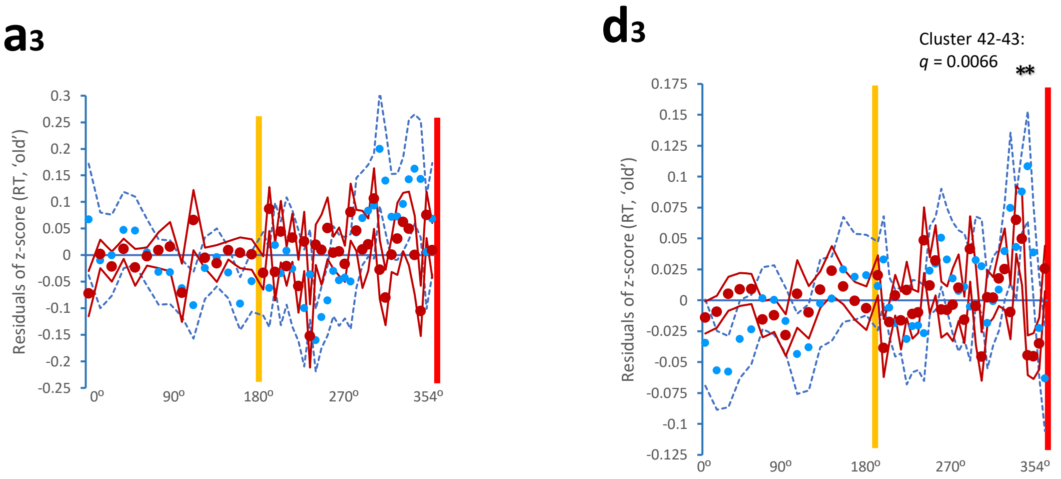


**Fig. S10. Re-binned analysis of cluster formations in RT of respiratory phase matching between sample and test blocks (1)**

Plots showing z-scored RTs (**a$_1$**, **b$_1$**, **c$_1$**, **D$_1$**), t-stats (**A$_2$**, **B$_2$**, **C$_2$**, **D$_2$**), and residuals of z-scored RTs (**A$_3$**, **B$_3$**, **D$_3$**) of 'old' events during the test respiratory cycle, according to four sample respiratory conditions: NoI$_s$ condition (**A$_1$-A$_3$**), IEt$_s$ condition (**B$_1$-B$_3$**), NoE$_s$ condition (**C$_1$, C$_2$**), and EIt$_s$ condition (**D$_1$-D$_3$**). We found skewed distributions of z-scored RT of the four sample respiratory conditions (*ps* ≤ 0.05, two-tailed one sample *t*-test against zero). The nonparametric permutation test and FDR correction revealed the significance of cluster formation (more than two adjacent bins) in the RT of NoI$_s$ condition (cluster 24-25: $t_{(sum)}$ = 4.87, *q* = 0.0010), and EIt$_s$ condition (cluster 40-41: $t_{(sum)}$ = 5.43, *q* = 0.00032). Residuals formed clusters in RT of EIt$_s$ condition (cluster 42-43: $t_{(sum)}$ = 4.60, *q* = 0.0066). * *p* ≤ 0.05 in **A$_1$**, **B$_1$**, **C$_1$**, and **D$_1$**. ** *q* ≤ 0.05 (a single cluster with two bins).

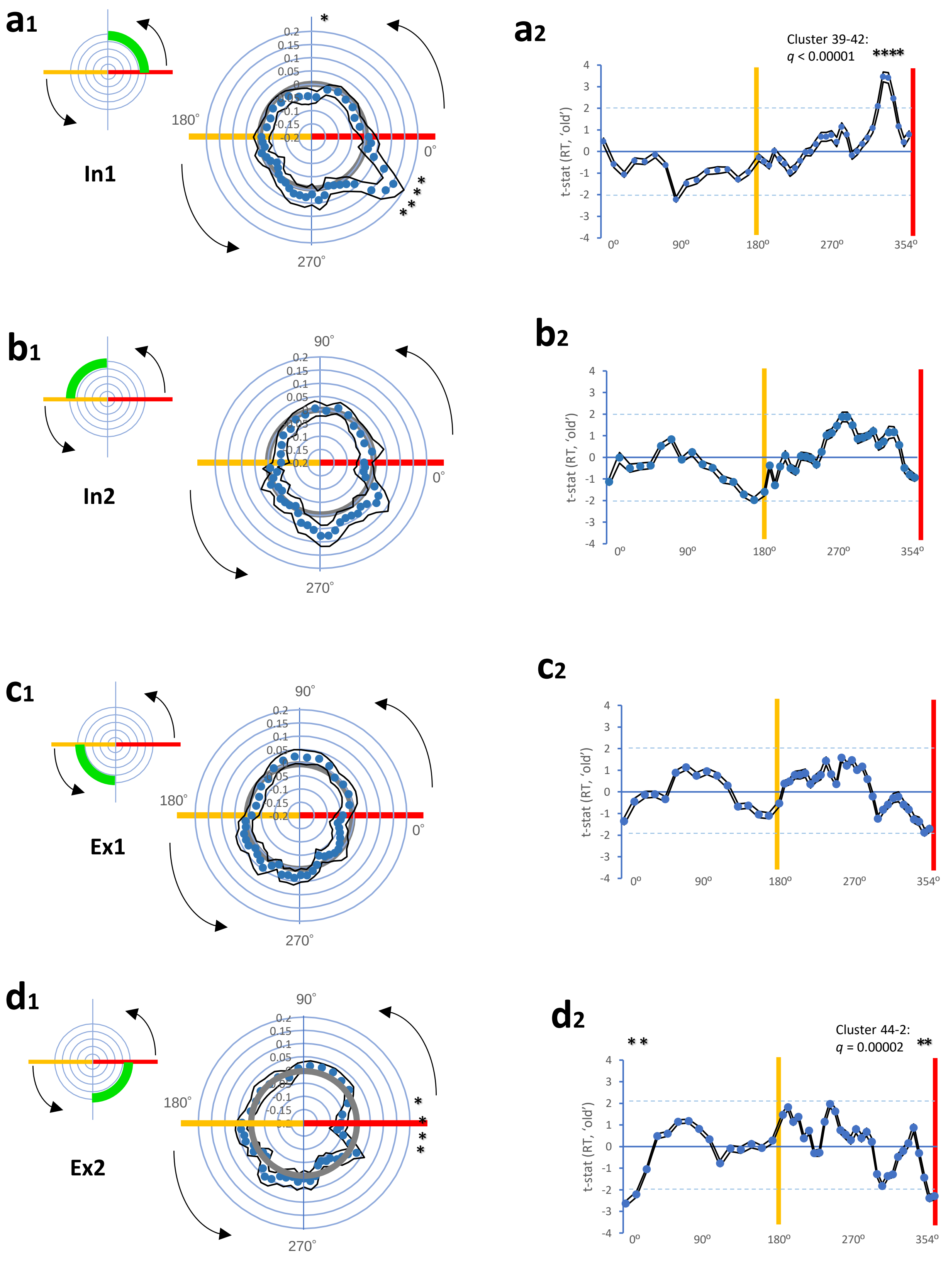

a1
In1
180°
0°
270°
a2
Cluster 39-42:
q < 0.00001
****
t-stat (RT, 'old')
0º
90º
180º
270º
354º
b1
In2
90°
0°
270°
b2
c1
Ex1
c2
d1
Ex2
d2
Cluster 44-2:
q = 0.00002
**

**a3**

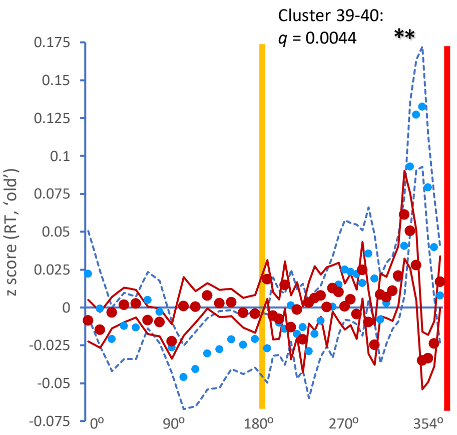


**d3**

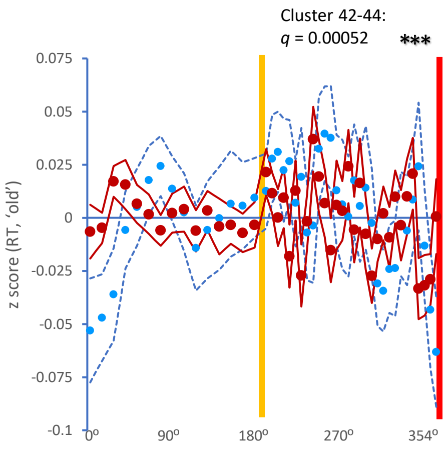


**Fig. S11. Re-binned analysis of cluster formations in RT of respiratory phase matching between sample and test blocks (2)**

Plots showing z-scored RTs (**$a_1$**, **$b_1$**, **$c_1$**, **$d_1$**), t-stats (**$a_2$**, **$b_2$**, **$c_2$**, **$d_2$**), and residuals of z-scored RTs (**$a_3$**, **$d_3$**) of 'old' events during the test respiratory cycle, according to four sample respiratory conditions: the sample In1 condition (**$a_1$-$a_3$**), sample In2 condition (**$b_1$, $b_2$**), sample Ex1 condition (**$c_1$, $c_2$**), and sample Ex2 condition (**$d_1$-$d_3$**). We found skewed distributions of z-scored RT of the sample In1 and sample Ex2 conditions (*ps* ≤ 0.05, two-tailed one sample *t*-test against zero). The nonparametric permutation test and FDR correction revealed the significance of cluster formation (more than two adjacent bins) in the RT of the sample In1 condition (cluster 39-42: $t_{(sum)} = 11.40$, $q < 1.0 \times 10^{-5}$), and the sample Ex2 condition (cluster 44-2: $t_{(sum)} = 9.55$, $q = 2.0 \times 10^{-5}$). Residuals formed clusters in RT of the sample In1 condition (cluster 39-40: $t_{(sum)} = 4.43$, $q = 0.0044$) and of the sample Ex2 condition (cluster 42-44: $t_{(sum)} = 7.05$, $q = 0.00052$). * $p \leq 0.05$ in **$a_1$** and **$d_1$**. ** $q \leq 0.05$ (a single cluster with two bins), *** $q \leq 0.05$ (a single cluster with three bins), **** $q \leq 0.05$ (a single cluster with four bins).
